# Pattern and Precision: DNA-Based Mapping of Spatial Rules for T Cell Activation

**DOI:** 10.1101/2025.06.12.659249

**Authors:** Shujie Li, Kaltrina Paloja, Maartje M.C. Bastings

## Abstract

The nanoscale spatial arrangement of T cell receptors (TCRs) ligands critically influences their activation potential in CD8^+^ T cells, yet a comprehensive understanding of the molecular landscape induced by engagement with native peptide–MHC class I (pMHC-I) remains incomplete. Using DNA origami as platform for multivalent engineering, we precisely organize pMHC-I molecules into defined spatial configurations to systematically investigate the roles of valency, inter-ligand spacing, geometric pattern, and molecular flexibility in regulating T cell function. We find that reducing inter-ligand spacing to ∼7.5 nm enhances T cell activation by up to eightfold compared to wider spacing (∼22.5 nm), and that as few as six pMHC-I molecules are sufficient to elicit a robust response. Notably, the geometry of pMHC-I presentation emerges as a key determinant of signaling strength, with hexagonal arrangements proving most effective. In contrast, the introduction of flexible linkers into pMHC-I impairs TCR triggering. Together, these findings define spatial parameters that govern pMHC-I–TCR interactions at the T cell interface and provide design principles for engineering next-generation T cell–based immunotherapies.

## INTRODUCTION

Cytotoxic T cell–mediated killing is a central mechanism for eliminating malignant cells and underpins the therapeutic success of adoptive T cell immunotherapy and immune checkpoint blockade. Activation of naïve CD8^+^ T cells begins with the engagement of the T cell receptor (TCR) by peptide–MHC class I (pMHC-I) complexes presented on the surface of antigen-presenting cells (APCs), such as dendritic cells (DCs). This communication between T cells and DCs is orchestrated by a diverse array of immune receptors whose spatiotemporal organization at the immunological synapse critically shapes the strength and quality of the T cell response^1^. Upon recognition of pMHC on DCs, TCRs rapidly organize into nanoclusters at the center of the T cell–DC interface, typically containing 5–30 receptors within a 35–70 nm radius^2^. This nanoscale organization has been shown to play a key role in modulating TCR signaling and likely reflects the spatial constraints required of its ligands^3^. As such, understanding the precise nanoscale requirements of TCR ligands that govern TCR triggering and T cell activation is essential and holds promise for the rational design of immune-modulating materials. However, mapping this molecular landscape within the complex context of T cell immunity remains a major challenge, as it demands platforms capable of controlling ligand presentation with nanometer-level precision.

The combination of high programmability and site-specific addressability endows DNA nanotechnology with a unique ability to control the spatial distribution of molecular components with nanometer precision^4,5^. As a result, functional DNA origami has enabled new paradigms for probing immunoreceptor–ligand interactions and holds significant promise for applications in cell-based immunotherapy^6–11^. Using DNA origami to present TCR-stimulating ligands such as anti-CD3ε antibodies (aCD3ε), previous studies have shown that CD8^+^ T cell activation reaches a plateau when three aCD3ε molecules are displayed, and increases as the inter-ligand spacing is reduced from 95 nm to 16 nm^12^. Similarly, in chimeric antigen receptor (CAR) T cell systems, optimal activation was achieved with four high-affinity DNA ligands arranged on DNA origami nanogrids, whereas further increasing ligand density resulted in premature termination of TCR signaling^13^. These findings, based on high-affinity TCR surrogates, provide valuable insights into the spatial constraints required for productive T cell activation. However, TCR triggering by high-affinity antibodies differs fundamentally from activation via the native ligand, pMHC^14^. As such, using pMHC is physiologically the most relevant for dissecting the spatial parameters underlying the local molecular landscape TCR activation.

More recently, the functional impact of pMHC spatial organization was investigated by manipulating two key parameters using DNA origami nanoscaffolds: density and inter-ligand spacing. Notably, densely clustered pMHC assemblies containing at least six streptavidin (SA) binding sites elicited stronger CD8^+^ T cell activation compared to free pMHC, with closer proximity between ligands correlating with enhanced activation^15^. While these findings underscore the importance of multivalent pMHC arrangement, the study did not fully disentangle the individual contributions of valency and density, as inter-ligand spacing was not uniformly controlled across different DNA origami designs. Additionally, pMHC clustering on triangular DNA origami was achieved through biotin–streptavidin interactions, where each tetravalent SA molecule could bind up to three biotinylated pMHC ligands^16^. This inherent multivalency complicates precise interpretation of how individual pMHC molecules contribute to T cell activation and obscures accurate measurement of true inter-ligand spacing.

While increasing attention has been given to ligand valency and inter-ligand spacing, the importance of local geometric pattern remains underexplored in the context of immune activation. We have previously shown that the geometric arrangement of multivalent ligands can critically determine the onset of super-selective receptor binding^17^, a phenomenon particularly relevant in the immune system, where ligand–receptor interactions are tightly regulated to preserve functional specificity and avoid aberrant activation. Indeed, CD95L arranged in a hexagonal pattern with 10 nm spacing on DNA origami was found to strongly induce CD95-mediated apoptotic signaling in activated immune cells, aligning precisely with the topography of CD95 receptor clusters at the cell surface^18,19^. Similarly, IgG antibodies have been shown to efficiently activate the complement system through ordered clustering into hexamers^20^. Moreover, the spatial patterning of PD-1 has been reported to influence its interaction with PD-L1 on dendritic cells, further supporting geometric organization as a key determinant of receptor engagement^21^. Beyond patterning, the local flexibility of ligand linkers is another critical design variable. Maintaining rigidity is essential for discerning spatial effects on immune receptor triggering. This principle has been demonstrated in studies involving the B cell receptor, Toll-like receptors, and Fas (CD95) signaling, where rigid presentation enhanced receptor activation^22–25^. Likewise, effector CD4^+^ T cells produce more cytokines in response to increasingly rigid substrates presenting TCR ligands, suggesting that mechanical properties of the interface also modulate downstream immune responses^26^. Despite growing interest, the combined impact of geometric pattern, rigidity, and linker flexibility on T cell signaling remains poorly characterized, representing a significant gap in our understanding of the spatial requirements for productive TCR triggering at the immunological synapse.

In this study, we systematically dissect how valency, inter-ligand spacing, geometric pattern, and ligand rigidity of pMHC-I independently influence T cell activation. By maintaining a constant inter-ligand spacing, we first investigated the effect of varying pMHC-I valency and found that multivalency linearly enhanced T cell activation at a spacing of ∼15 nm. Reducing the inter-ligand distance to ∼7.5 nm further amplified the response, with just six pMHC-I molecules sufficient to elicit robust T cell activation. We next explored the roles of geometric pattern and molecular flexibility, demonstrating that a compact hexagonal arrangement of pMHC-I elicited the strongest activation, while ligand flexibility impaired TCR signaling by disrupting the tight spatial constraints required for productive TCR–pMHC engagement. Together, our findings provide new insights into the nanoscale spatial requirements governing pMHC– TCR interactions and underscore the importance of precisely tailoring spatial parameters in the rational design of multivalent materials for T cell–based immunotherapies.

## RESULTS AND DISCUSSION

### Dense pMHC-I Clusters Reveal a Spatial Threshold for Efficient T Cell Stimulation

To investigate how TCR ligand valency and inter-ligand spacing influence T cell activation, we used a two-layer, 60 nm diameter disk-shaped DNA origami nanoplatform. This structure was strategically chosen to mimic the geometry of the immunological synapse and offers robust site-specific positioning of macromolecules^21^ **(Figure S1)**. We generated a library of DNA disks presenting ovalbumin (OVA_257–264_; SIINFEKL)-loaded MHC-I molecules (termed pMHC-I) with controlled valencies (0, 4, 6, 8, and 12) and inter-ligand spacings (7.5 nm or 15 nm), enabling a systematic evaluation of their individual and combined effects on the activation of primary CD8^+^ T cells **(Figure 1a, b; Figure S2, S3)**. At 7.5 nm spacing, T cell activation increased with rising pMHC-I valency from 0 to 6, confirming a positive correlation between ligand number and TCR triggering as reported previously^15,27^ **(Figure 1c)**. However, increasing the valency beyond six yielded minimal additional benefits in terms of activation markers CD69, CD25, and CD137, which represent early, late, and very late activation, respectively. While a slight resurgence in activation was observed at valency 12, this did not translate into increased cytotoxic function, as measured by expression of FasL (a death-inducing ligand) and CD107a (a marker of degranulation). Cytokine secretion levels, including TNF-α and IFN-γ, aligned with the surface marker data, further supporting this saturation effect **(Figure 1c)**. When the inter-ligand spacing was increased from 7.5 nm to 15 nm, T cell activation decreased across all valency conditions, although the extent varied. Notably, for six pMHC-I ligands, CD69 expression dropped sharply at the wider spacing, whereas constructs presenting four ligands showed a more modest decline. Interestingly, at 15 nm spacing, valency became the dominant determinant of T cell activation, with activation levels increasing approximately linearly from valency 0 to 12 **(Figure 1c)**. This suggests that tight spatial organization is particularly important at lower ligand densities, while higher valency can partially compensate for suboptimal spacing.

**Figure 1.**
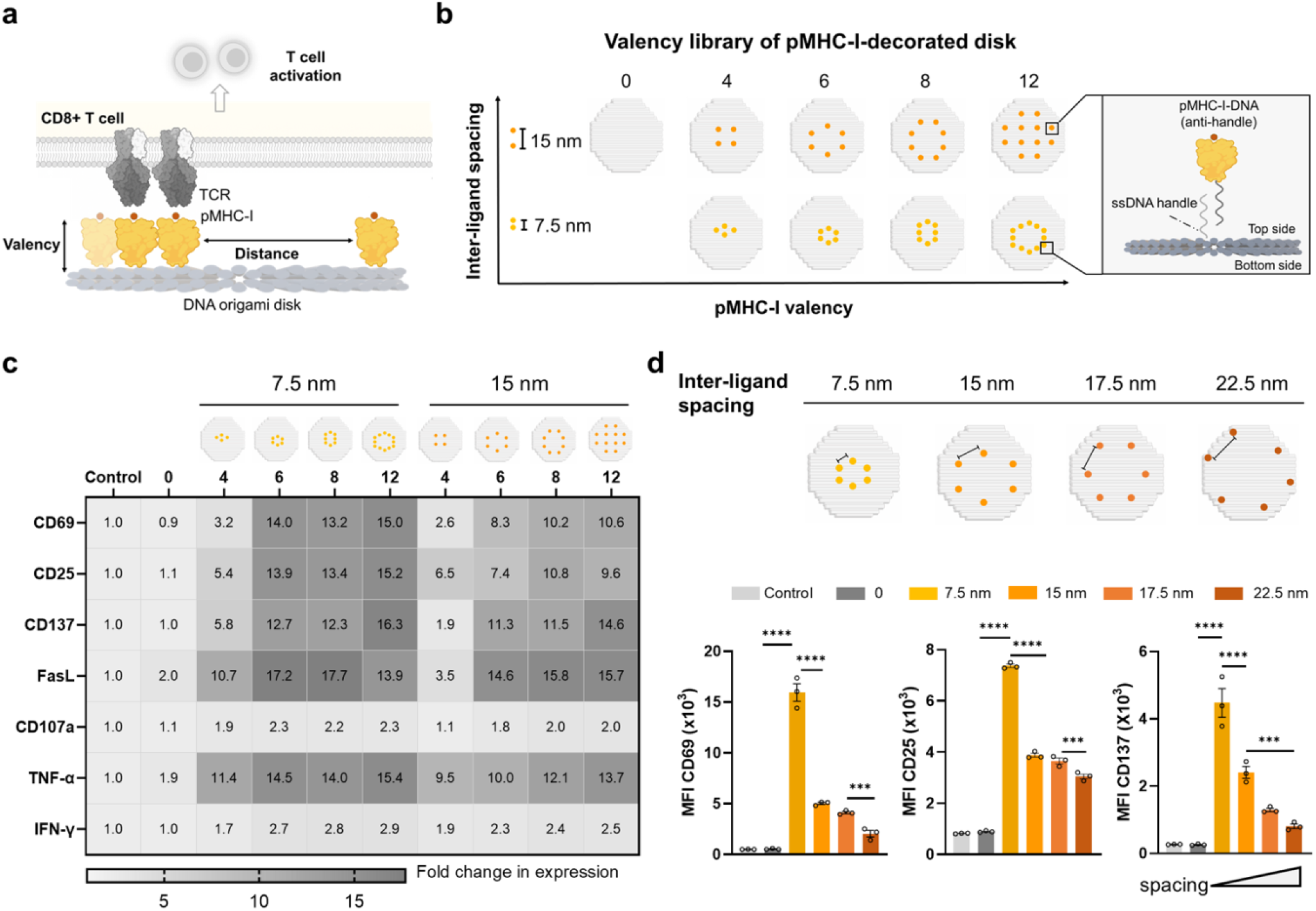
Effects of pMHC-I valency and inter-ligand spacing on CD8^+^ T cell activation. (a) Schematic presentation of exploring the impact of clustering differences (valency and inter-ligand spacing) of pMHC-I on CD8^+^ T cell activation with DNA origami disk nanoplatform; (b) Schematic presentation of the DNA origami disk designs used in the T cell activation assay. Engineered pMHC-I was decorated onto the DNA disk via handle-anti-handle hybridization; (c) Flow cytometry analysis of biomarkers and cytokine expression in CD8^+^ T cells following stimulation with DNA disks at 24 h (CD69), 48 h (CD25, TNF-α, IFN-γ), and 72 h (CD137, FasL, CD107a). Fold changes in expression are represented as the mean fluorescence intensity (MFI) values relative to the untreated control group (n≥6 from two independent experiments); (d) MFI of CD69 (24 h), CD25 (48 h), and CD137 (72 h) in CD8^+^ T cells stimulated by DNA origami presenting pMHC-I with varied inter-ligand spacing. Data are shown as mean ± SEM (n=3 biological replicates). Statistical significance was assessed using one-way ANOVA with Tukey’s multiple comparisons test (\*\*\**p* < 0.001, \*\*\*\**p* < 0.0001).

We hypothesized that further reducing the inter-ligand spacing, particularly when using the physiological TCR ligand pMHC, might enhance T cell activation by better matching native TCR spatial constraints. This hypothesis was supported by our data, which revealed a clear distance-dependent trend in T cell activation using pMHC-I disks at a fixed valency of six: 7.5 nm > 15 nm > 17.5 nm > 22.5 nm **(Figure 1d; Figure S4)**. These results suggest that tighter spatial organization of ligands facilitates more efficient TCR triggering. This finding is consistent with prior results showing that three aCD3 antibodies spaced at 16 nm can induce saturated T cell activation^12^. As antibodies engage with 2 binding epitopes, effectively 6 interactions in 8 nm distance are formed. The robust activation we observed with six pMHC-I molecules spaced at 7.5 nm indicates that this distance may approximate a lower functional limit for effective ligand spacing on synthetic platforms. Taken together, our results demonstrate that a minimal unit of six pMHC-I molecules arranged at 7.5 nm spacing is sufficient to induce strong T cell activation. This finding highlight that a low copy number of locally clustered native ligands is enough to stimulate a potent T cell response^28^.

### Hexagonal ligand patterning maximizes T cell activation

To investigate whether the spatial patterning of pMHC-I molecules influences T cell activation, we assembled DNA disks presenting six pMHC-I ligands arranged in hexagonal (Hex), triangular (Tri), linear (Lin), and parallel (Par) configurations. Based on the consensus from our previous experiments, the ligand copy number and inter-ligand spacing were fixed at six and 7.5 nm, respectively **(Figure 2a; Figure S5a, b)**. Strikingly, we observed that the geometric pattern of pMHC-I had a significant impact on both CD8^+^ T cell activation and cytotoxic function **(Figure 2b; Figure S5c)**. In particular, the linear configuration resulted in substantially lower expression of activation markers CD69, CD25, and CD137, as well as effector molecules FasL and CD107a, compared to the other patterns **(Figure 2b; Figure S5c)**. This reduction is likely due to the mismatch between the linear arrangement of ligands and the clustered distribution of TCRs on the T cell surface, which favors engagement by locally clustered ligands^29,30^.

**Figure 2.**
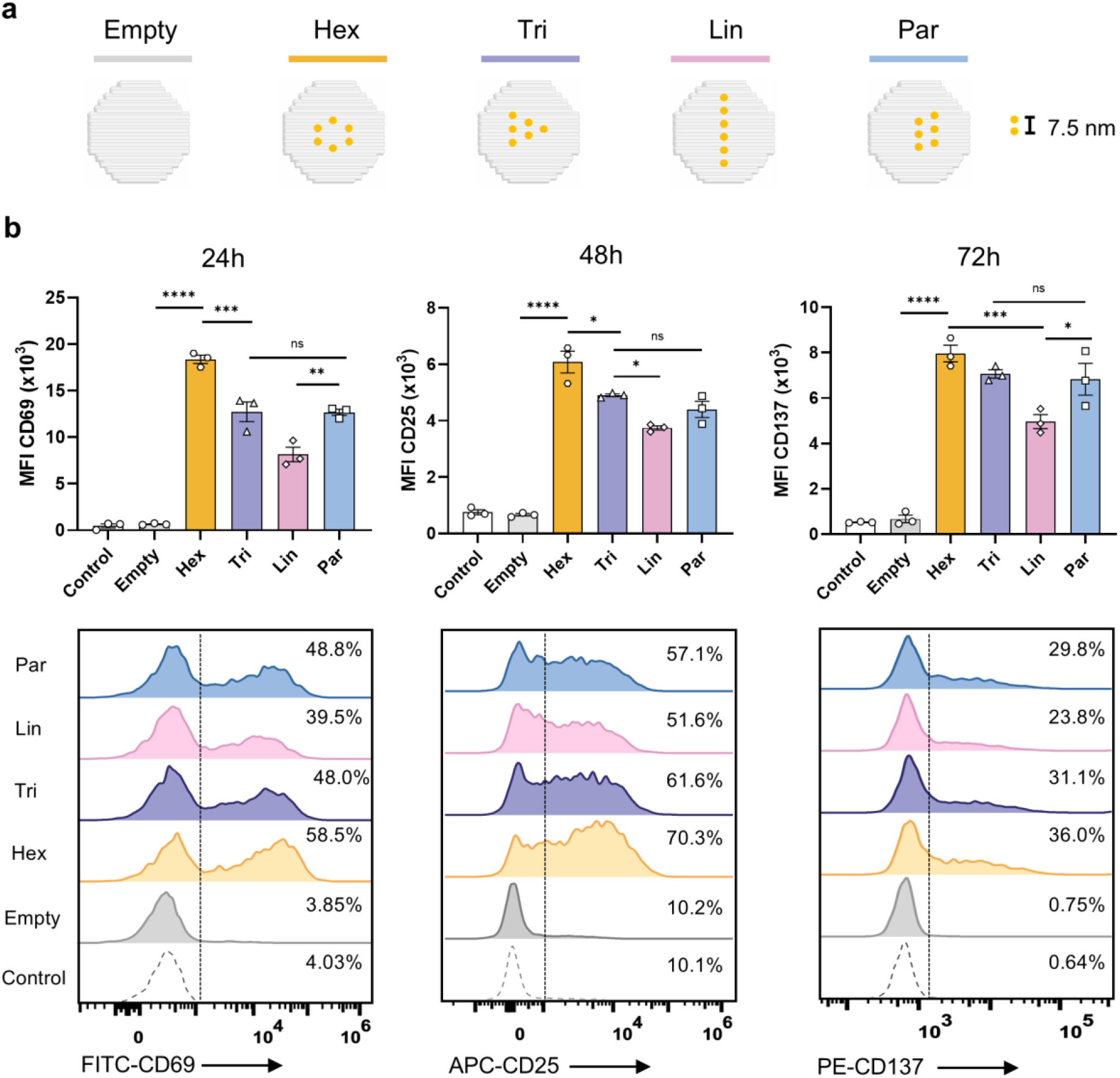
Effects of pMHC-I geometric pattern on CD8^+^ T cell activation. (a) Schematic presentation of DNA origami disk displaying pMHC-I in four different geometric patterns. Hex: hexagonal, Tri: triangular, Lin: linear, Par: parallel. The inter-ligand spacing of pMHC-I was 7.5 nm. (b) Expression levels of CD69, CD25, and CD137 on CD8^+^ T cells (upper panel) and representative flow cytometry data (lower panel) following stimulation with pMHC-I-patterned DNA disks, showing T cell activation at 24, 48, and 72 hours, respectively. Percentage of CD69+, CD25+, and CD137+ cells of each group were gated based on untreated control group. Data are shown as mean ± SEM (n=3 biological replicates). Statistical significance was assessed using one-way ANOVA with Tukey’s multiple comparisons test (\**p* < 0.05, \*\**p* < 0.01, \*\*\**p* < 0.001, \*\*\*\**p* < 0.0001).

DNA disks presenting pMHC-I in triangular and parallel patterns elicited comparable levels of T cell activation, suggesting that local clustering, regardless of precise pattern, supports efficient TCR triggering. However, the hexagonal arrangement induced the strongest activation profile, with 58.5% CD69^+^, 70.3% CD25^+^, and 36% CD137^+^ T cells, substantially higher than both the triangular and parallel configurations **(Figure 2b)**. One possible explanation is that the higher ligand density in the parallel and triangle patterns may cause steric hindrance, limiting full TCR engagement **(Figure S5d)**. Alternatively, the hexagonal arrangement, which has been reported to optimize antibody binding^31^, may provide greater geometric symmetry and multidirectional accessibility, thereby increasing the likelihood of TCR engagement, promoting stabilization of TCR nanoclusters, and facilitating the recruitment of downstream signaling molecules.

At the population level, the hexagonal configuration maximized expression of all monitored biomarkers. The mean fluorescence intensity (MFI) for CD69 was 2.3-fold higher, and for CD25 and CD137 was 1.6-fold higher, compared to the linear pattern **(Figure 2b)**. Importantly, these measurements were obtained in live cells, where TCRs can laterally diffuse and reorganize. Despite this membrane fluidity, the pattern-dependent differences in activation remained robust and detectable. While previous studies have predominantly emphasized the role of ligand density in TCR triggering^15^, our findings highlight the critical and often overlooked role of geometric pattern, a spatial design parameter that may play a key role in natural immune cell–cell communication mediated by multivalent ligand–receptor interactions.

### Flexibility Compromises Geometry-Driven TCR Triggering

Linker flexibility plays a key role in the design of functional nanomaterials by shaping the spatial interface through which ligands engage cell surface receptors. This parameter is especially critical in immune signaling, where the immunological synapse is both spatially and temporally confined. Prior studies have shown that immune synapse formation is most dynamic within the first 10 minutes of T cell–APC contact and remains detectable up to an hour later, indicating that TCR clusters exhibit constrained, directed motion during early activation^32^. This suggests that receptor engagement may require a certain degree of positional rigidity. In support of this, extending the linker length between pMHC dimers and thereby introducing flexibility, has been shown to disrupt TCR triggering^33,34^. Motivated by these insights, we examined the influence of ligand flexibility on CD8^+^ T cell activation by modifying the rigidity of the DNA–pMHC-I interface. Specifically, we incorporated thymine (T) spacers into the ssDNA handle extending from the DNA disk, presenting pMHC-I ligands with increasing degrees of flexibility. We focused on the hexagonal (Hex_0T, Hex_5T, Hex_10T) and linear (Lin_0T, Lin_5T, Lin_10T) configurations, which previously showed the strongest and weakest activation potentials, respectively **(Figure 3a, c; Figure S6a, b)**.

**Figure 3.**
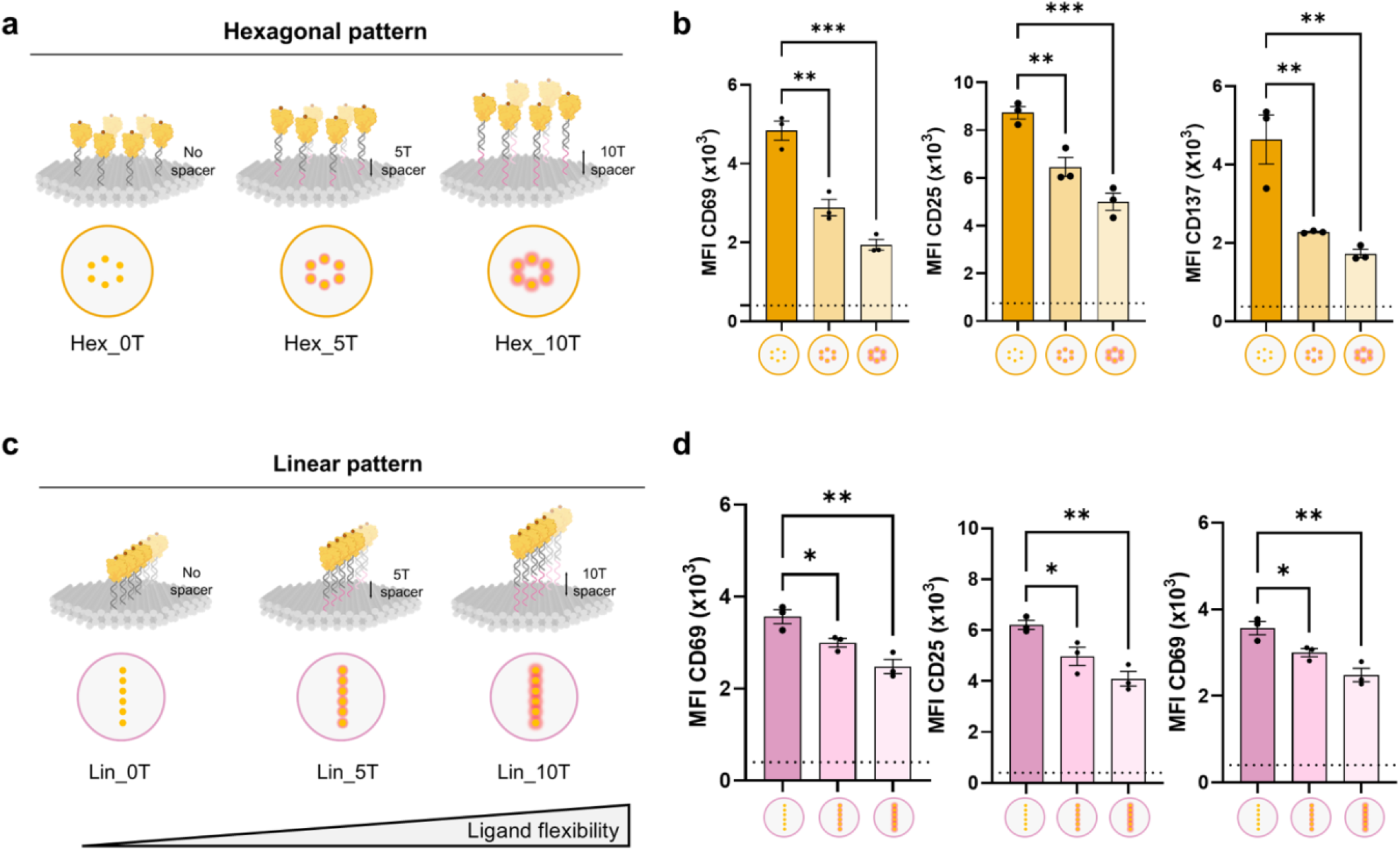
Effects of pMHC-I ligand flexibility on CD8^+^ T cell activation. (a) Schematic illustration of DNA origami disks presenting pMHC-I with increasing ligand flexibilities arranged in a hexagonal pattern. (b) Expression levels of CD69 (24 h), CD25 (48 h), and CD137 (72 h) on CD8^+^ T cells stimulated by hexagonally patterned pMHC-I DNA origami with varying ligand flexibilities. (c) Schematic illustration of DNA origami disks presenting pMHC-I with increasing ligand flexibilities arranged in a linear pattern. (d) Expression levels of CD69 (24 h), CD25 (48 h), and CD137 (72 h) on CD8^+^ T cells stimulated by linearly patterned pMHC-I DNA origami with varying ligand flexibilities. The dashed line indicates the corresponding biomarker expression induced by empty DNA origami disk without pMHC-I decoration. Data are shown as mean ± SEM (n=3 biological replicates). Statistical significance was assessed using one-way ANOVA with Tukey’s multiple comparisons test (\**p* < 0.05, \*\**p* < 0.01, \*\*\**p* < 0.001).

Interestingly, the rigid Hex_0T configuration elicited significantly higher expression of activation markers CD69, CD25, and CD137 at both early and late time points compared to its more flexible counterparts (Hex_5T and Hex_10T) **(Figure 3b; Figure S6c)**. T cell activation was progressively dampened as linker flexibility increased, with a consistent trend observed across both hexagonal and linear patterns **(Figure 3b, d; Figure S6d)**, indicating a negative correlation between ligand flexibility and T cell activation. Importantly, the geometric pattern effect remained evident: hexagonal configurations consistently outperformed linear ones in stimulating T cells. However, the magnitude of this difference diminished with increasing linker flexibility, suggesting that the mechanical instability introduced by flexible linkers compromises the benefits of geometric presentation **(Figure S7)**. Similar trends were also observed for the parallel configuration, reinforcing the general principle that rigidity enhances TCR triggering across diverse spatial patterns **(Figure S8)**.

### Mechanistic Insight into Ligand Flexibility and Spatial Tolerance

To more precisely understand how ligand flexibility modulates T cell activation, we examined the underlying mechanisms involved. Previous studies have shown that spatial tolerance, a combination of ligand rigidity and matched receptor spacing, is essential for functional interactions, such as CpG–TLR9 binding^24^. In our system, introducing 5T or 10T single-stranded DNA spacers into the ligand handle increases positional uncertainty by approximately ±1.7 nm or ±3.4 nm, respectively, assuming full extension **(Figure 4a)**. This spatial deviation likely impairs the alignment between pMHC-I and TCR, as reflected by the diminished T cell activation observed with more flexible constructs **(Figure 3)**. Given that T cell activation is initiated via TCR–pMHC engagement, which in turn activates a tyrosine kinase cascade, we hypothesized that ligand flexibility disrupts early TCR signaling. To test this, we monitored phosphorylation levels of ZAP70, ERK1/2, and NF-κB (p65), 15 minutes after stimulation. As expected, stimulation with non-functionalized DNA disks did not induce signaling above background levels seen in unstimulated CD8^+^ T cells **(Figure 4b)**, enabling a clear assessment of the effects of ligand flexibility. Consistent with activation marker expression, the highest phosphorylation levels of ZAP70, ERK1/2, and NF-κB were observed in cells treated with rigid pMHC-I ligands (0T spacers), regardless of geometric pattern (hexagonal or linear) **(Figure 4b)**. Intermediate signaling was observed for 5T constructs, while the 10T variants showed the weakest phosphorylation, further confirming that ligand flexibility negatively impacts TCR signaling efficiency.

**Figure 4.**
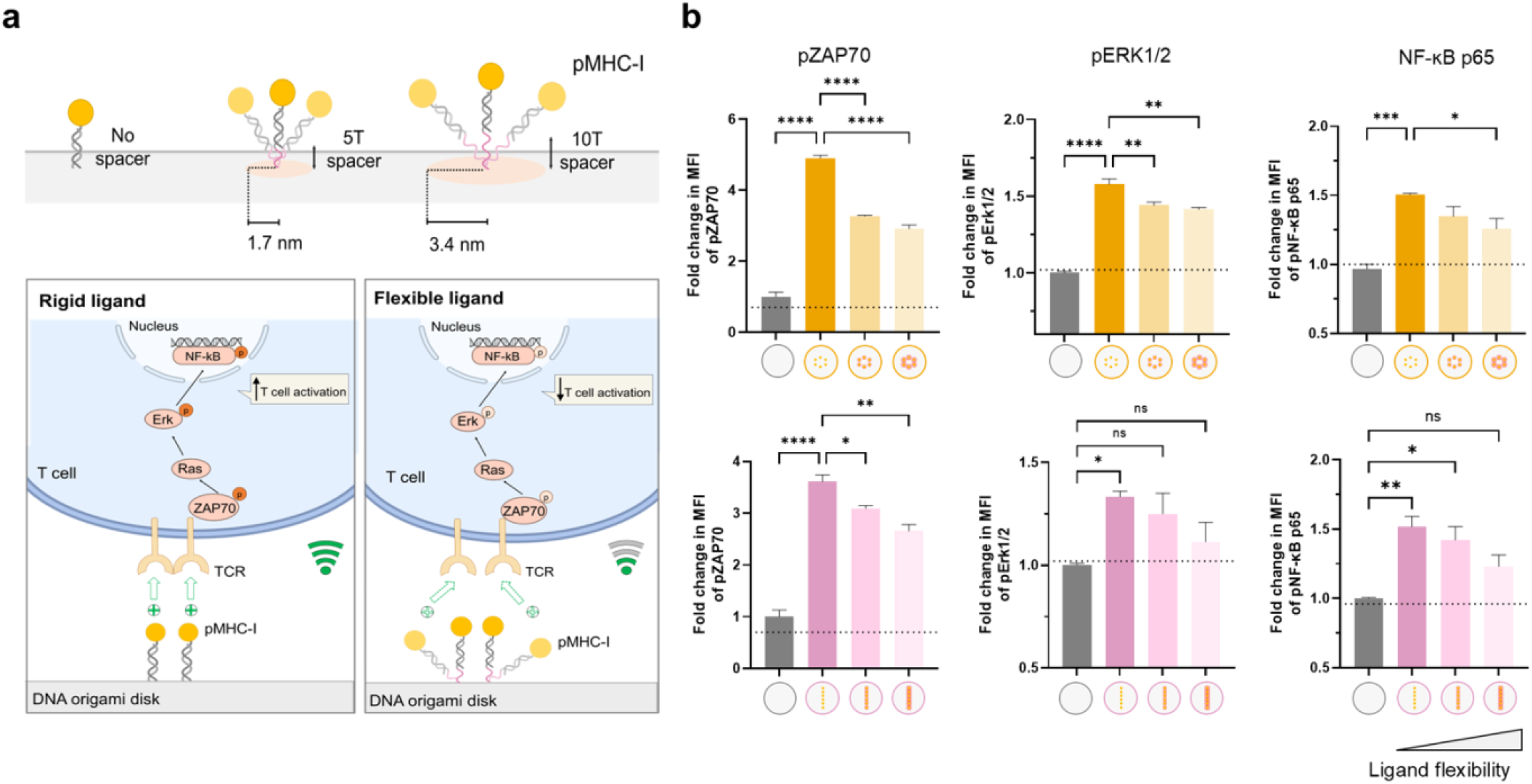
Effects of pMHC-I ligand flexibility on TCR triggering. (a) Schematic representation of the estimated spatial deviations in pMHC-I positioning induced by 5T or 10T linkers (upper panel), and the corresponding TCR signal transduction triggered by rigid versus flexible pMHC-I ligands (lower panel). (b) Relative MFI of phosphorylated ZAP70, ERK1/2, and NF-κB (p65) after 15 min stimulation with pMHC-I arranged in hexagonal (upper panel) or linear (lower panel) patterns on DNA origami disk, with varying ligand flexibility. Data were normalized to the response from empty DNA origami without pMHC-I decoration. The dashed line indicates the protein phosphorylation level in unstimulated CD8^+^ T cells. Data are shown as mean ± SEM (n=3 biological replicates). Statistical significance was assessed using one-way ANOVA with Tukey’s multiple comparisons test (\**p* < 0.05, \*\**p* < 0.01, \*\*\**p* < 0.001, \*\*\*\**p* < 0.001).

We conclude that increased ligand flexibility disrupts the nanoscale coordination between pMHC-I and TCRs by introducing spatial deviations that reduce the frequency of productive short-distance TCR–pMHC interactions. To validate this, we compared flexible ligands (Hex_5T, Hex_10T) to rigid pMHC-I ligands spaced at 15 nm and 22.5 nm, respectively. As shown in **Figure S9**, the activation potency of flexible ligands fell between the responses induced at 7.5 nm and 15 nm spacing, consistent with predicted effects of spatial mismatch. Importantly, TCR triggering is highly sensitive to spatial perturbations at the immune synapse, not only because of ligand–receptor geometry but also due to the biophysical organization of the signaling interface. According to the kinetic segregation model, TCR signaling is initiated when close-contact zones between the T cell and antigen-presenting cell exclude large membrane phosphatases such as CD45, whose bulky extracellular domain (∼30–50 nm) sterically prevents access to regions where the membranes are tightly apposed^35–37^. This exclusion allows kinases to phosphorylate ITAM motifs in the CD3 complex, thereby initiating downstream signaling. Given that the extra distance due to the T-spacers is <5nm, we consider the effects of CD45 not important. Taken together, our results reveal a spatial tolerance window for productive TCR triggering, demonstrating that even nanometer-scale deviations in ligand positioning, whether from increased spacing or flexibility, can significantly affect immune signaling. These findings underscore the extreme spatial sensitivity of the TCR–pMHC interface and reinforce the need for rigid, geometry-controlled presentation of ligands in the design of immunomodulatory materials.

## Conclusion

While extensive efforts have been directed toward developing immune-engineering biomaterials to support T cell expansion and adoptive cell therapies, a comprehensive understanding of how the spatial presentation of TCR ligands governs T cell activation remains limited. Leveraging the unparalleled spatial precision of DNA nanotechnology, we systematically dissected the functional impact of four key parameters of native TCR ligand (pMHC-I) presentation: valency, inter-ligand spacing, geometric pattern, and ligand rigidity. By monitoring CD8^+^ T cell responses, we found that short inter-ligand spacing (7.5 nm) substantially enhances activation, and that as few as six pMHC-I molecules, when locally clustered, are sufficient to trigger a robust response. Notably, we show that ligand geometry plays a decisive role: hexagonally arranged pMHC-I outperforms linearly arranged ligands of identical valency, highlighting the importance of matching ligand spatial organization to receptor topology on the T cell surface. Furthermore, we demonstrate that ligand flexibility impairs TCR engagement and signaling, likely by disrupting the membrane proximity and nanoscale clustering required for productive TCR–pMHC interaction and CD45 exclusion.

Together, these findings emphasize that effective multivalent interactions rely not only on ligand density, but on the precise engineering of spatial parameters including pattern symmetry and mechanical stability. This represents a conceptual shift from conventional multivalent binding to what we define as multivalent engineering: the deliberate spatial programming of ligand organization to engage biological interfaces with maximal functional control. In summary, our study delineates the spatial principles governing receptor–ligand interactions at the T cell interface and establishes a framework for the rational design of immunotherapeutic materials. By tuning valency, spacing, pattern, and rigidity, multivalent systems can be tailored to more effectively coordinate receptor engagement and downstream signaling, paving the way toward next-generation materials for T cell–based immunotherapy.

## Acknowledgments

We are grateful for the Protein Production and Structure Core Facility (PTPSP, EPFL) and the Flow Cytometry Core Facility (FCCF, EPFL) for instrument accessibility and operational support, and the Center of PhenoGenomics (CPG, EPFL) for housing the animals. The authors thank Yameng Lou and Dr. Vincenzo Caroprese for their insightful discussions.

