## Supporting Info for "Pattern and Precision: DNA-Based Mapping of Spatial Rules for T Cell Activation"

### METHODS AND MATERIALS

**Preparation of bare DNA origami disks.** Bare DNA disks were assembled as previously described[1]. To stabilize molecular arrangement and positioning to ensure effective and precise TCR-MHC binding, DNA disks were biotinylated on the bottom side for the purpose of immobilization onto a streptavidin-coated surface **(Figure S1)**. Briefly, a mixture pool consists of 10 nM of ssDNA scaffold p7560 (Tilibit nanosystems) and 10x/5x/7x molar excess of unmodified DNA staple strands/Cy5-integrated DNA strands/biotin-integrated DNA strands (IDT) was prepared in folding buffer (5 mM Tris pH 8, 1 mM EDTA, 5 mM NaCl, 18 mM $MgCl_2$). The mixture was subjected to an annealing program started with 5 min at 80°C, then cooled from 60°C to 20°C in steps of −1°C per hour in a thermal cycler (Biometra TRIO). Assembled structures were purified by adding an equal volume of PEG precipitation buffer (15% PEG8000 (w/v), 5 mM Tris, 1 mM EDTA, 500 mM NaCl, 18 mM $MgCl_2$) to the reaction solution. After centrifugation at 16,000 × *g* for 40 min at 20°C, the purified DNA disk pellets were collected and resuspended in 10 mM HEPES pH 7.4, 150 mM NaCl supplemented with 18 mM $MgCl_2$. The concentration of bare DNA disks was determined by measuring absorption at 260 nm on a Nanodrop (Q9000, Quawell). Structure folding was verified by agarose gel electrophoresis (1% agarose (w/v), 0.5x TBE buffer (B52, Thermo Scientific), 18 mM $MgCl_2$, and 1x SYBR Safe) conducted at 70 V for 90 min on ice. Gels were imaged with a Chemidoc MP imaging system. The bare DNA disk samples were stored at 4°C until further use.

**Production of single-chain SIINFEK-bound MHC-I (H-2Kb).** The gene encoding for mouse SIINFEKL MHC-I was constructed from a published sequence the following modifications. Specifically, the sequence was designed with a C-terminus sortase A (StrA) recognition motif LPETG as a functionalization site, followed by an 8x-His tag for purification purpose. A BiP signal peptide was modified at the N-terminus for protein secretion. The detailed construct design and protein sequence have been described in a previously publication from our laboratory[2]. SIINFEKL MHC-I (H-2Kb) was expressed in insect cells using Bac-to-Bac baculovirus expression system. Briefly, the engineered MHC-I gene was cloned under polyhedrin promoter into pFastBac1 vector (GenScript Biotech). Recombinant bacmid was prepared by transforming the plasmid into competent cells DH10EMBacY. Fresh Sf21 cells were transfected with the bacmid using transfection agent (XtremeGENETM 9; Sigma-Aldrich) to generate recombinant baculovirus (rBV). For MHC-I expression, $1x10^6$/mL Hi5 cells were infected with rBV for ~72 h at 28°C with agitation. The supernatant was collected and purified by Fastback Ni advance resin (Fastback-Ni-Adv-10, Protein Ark) and subsequent size exclusion chromatography (SEC) using a Superdex 200 Increase 10/300 GL column on ÄKTA systems. The purity of MHC-I protein was evaluated by SDS-PAGE analysis **(Figure S2a)**. The protein was stored at −80°C until further use.

**StrA-mediated MHC-I-DNA conjugation.** Purified pMHC-I was labeled with ssDNA strand via sortase-mediated reaction and copper-free click chemistry. Prior functionalization, sortase A5 (SrtA) was generated using the same method as previously described[3] **(Figure S2b)**. For DBCO ligation, 20 µM pMHC-I, 20 µM SrtA, and 2 mM amine-DBCO linker (761540, Sigma-Aldrich) were reacted in 20 mM HEPES pH 7.4, 150 mM NaCl supplemented with 10 mM $CaCl_2$ at 4°C. After 16 h incubation, excessive linker was removed by buffer exchange to 20 mM HEPES, 150 mM NaCl using 7K MWCO Zeba Spin Desalting columns (89882, Thermo Scientific). The pure pMHC-I-DBCO conjugates were obtained by removing His-tagged unreacted pMHC-I and SrtA through HisPur™ Ni-NTA resin (88221, Thermo Scientific). For DNA labeling, 4-fold molar excess of azide-ssDNA (anti-handle) (/5'azide/CATGTCAGGAGATTTTCAGCC, IDT) was added to pMHC-I-DBCO conjugate and incubated overnight at 4°C. The reaction mixture was purified with 30K MWCO Amicon centrifugal filters (UFC9030, Sigma-Aldrich) pre-coated with Pluronic F-127 (P2443, Sigma-Aldrich, 5% (w/v) in water) to remove excess azide-ssDNA. The concentration of pMHC-I-DNA conjugate was measured using a Micro BCA assay kit (23235, Thermo Scientific).

To further verify the presence of DNA on the conjugate, pMHC-I-DNA conjugate was incubated with 2.5-fold molar excess of Cy5-labeled complementary ssDNA strand (Cy5-compDNA, /5'Cy5/GGCTGAAAATCTCCTGACATG) for 1 h at 30°C. Samples were loaded on a 4-12% Tris-Glycine gel (XP04125BOX, Thermo Scientific). The gel was run at 200 V for 45 min and visualized with Chemidoc MP imaging system (Bio-Rad) **(Figure S2 b, c)**.

**Preparation of pMHC-I decorated DNA origami disks.** For pMHC-I attachment, the staples corresponding to the modification sites were extended at their 3′ end with a 21-nucleotide handle. For functionalization, purified bare DNA disks were incubated with 6-molar excess of pMHC-I-DNA conjugate per protruding handle for 1 h at 28°C followed by 14 h at 22°C in thermal cycler. Excess proteins were removed using 100K MWCO Amicon centrifugal filters (UFC510008, Sigma-Aldrich) pre-coated with 5% Pluronic F127 (w/v). Similarly, the concentration of pMHC-I DNA disks were quantified on nanodrop. 2% agarose gel (w/v) electrophoresis was performed to confirm the decoration of proteins on DNA disks. For better observation of band shift, the gel was run at 70 V for 140 min on ice. Purified pMHC-I decorated DNA disks were stored at 4°C and used within three days.
DNA sequences are available in **Tables S1-S7**.

**Animals.** TCR transgenic OT-I mice (C57BL/6-Tg(TcraTcrb)1100Mjb/J, female, 8- to 13-week-old) were purchased from Charles River (France) and maintained at the animal facility in the Center of PhenoGenomics (EPFL, Switzerland). Animal husbandry and euthanasia for the extraction of spleens and lymph nodes (LNs) were performed in compliance with protocols approved by the veterinary authorities of Canton de Vaud according to the Swiss regulations (license VD4004).

**T cell isolation.** Mouse spleen and axillary and inguinal LNs collected from OT-I mice were ground through a 40 μm cell strainer (352340, Corning). Afterwards, primary OT-I $CD8^+$ T cells were isolated from splenocytes and lymphocytes using the EasySep™ mouse $CD8^+$ T cell isolation kit (19853, Stemcell Technologies) according to manufacturer's instructions. Pure $CD8^+$ T cells were maintained in T cell medium (RPMI 1640-GlytaMAX medium (61870010, Thermo Scientific) supplemented with heat-inactivated 10% FBS (v/v), 1% penicillin/streptomycin (v/v), 10 mM HEPES, and 50 μM β-mercaptoethanol at 4°C until further use. To examine the purity of isolated $CD8^+$ T cells, cells were stained with LIVE/DEAD™ fixable violet dead cell stain (L34955, Thermo Scientific) for 30 min at 4°C, and PE anti-mouse CD3ε (500A2, BioLegend) and APC anti-mouse CD8a (53-6.7, BioLegend) antibody for 30 min at 4°C, followed by analysis on CytoFLEX flow cytometer (Beckman). The purity was typically 93-97% **(Figure S10a)**.

**Immobilization of DNA disks.** Prior to seeding $CD8^+$ T cells, high-binding 96-well plates (3361, Corning) were coated with streptavidin (300 nM in PBS, 100 μL per well) (21122, Thermo Scientific) overnight at 4°C, washed twice with PBS, and blocked with 3% BSA solution (in PBS, w/v) for 30 min at 37°C. Afterwards, corresponding biotinylated pMHC-I decorated DNA disks (8 nM, 35 μL per well) were added to the streptavidin-precoated plates and incubated on a shaker for 45 min at room temperature. Uncoated disks were removed by washing the plate twice with 10 mM HEPES pH 7.4, 150 mM NaCl, 18 mM $MgCl_2$. DNA disk immobilization level was monitored using Cy5 intensity measured with Cytation 5 plate reader (BioTek Instruments).

**Cytotoxicity assay.** To assess the cytotoxic effects of DNA origami disks and $MgCl_2$ supplementation on $CD8^+$ T cells, 50,000 $CD8^+$ T cells were seeded into empty DNA disk-immobilized plates and cultured in T cell medium supplemented with 15 mM $MgCl_2$. Control groups included untreated cells and cells treated with 15 mM $MgCl_2$ in the absence of DNA disks. After 24, 48, and 72 hours of co-incubation, cell viability was measured using the Cell Counting Kit-8 (HY-K0301, MedChemExpress) following the manufacturer's instructions **(Figure S10b)**.

**In vitro T cell activation assay.** $CD8^+$ T cells (50,000 per well) were seeded into the corresponding DNA disk-immobilized plates and cultured in T cell medium in the presence of 0.5 μg/mL soluble mouse anti-CD28 antibody (102116, BioLegend), 10 ng/ml recombinant mouse IL-2 (575402, BioLegend), 5 ng/ml recombinant mouse IL-7 (577802, BioLegend), and 15 mM $MgCl_2$ **(Figure S1b)**. The cells were incubated at 37°C with 5% $CO_2$ in a humidified incubator for the indicated durations. T cells were cultured for a total of 24 h, 48 h, or 72 h and every 24 h cells were transferred on freshly prepared DNA disk-immobilized plates.

For flow cytometry analysis, at the end of the treatment period, T cells underwent violet live/dead staining (L34955, Thermo Scientific) at 4°C for 30 min, and subsequent surface marker staining in BD Pharmingen stain buffer (BSA) (554657, BD Biosciences) with corresponding fluorescently labeled anti-mouse antibodies (FITC anti-CD69 (104505), APC anti-CD25 (102011), PE anti-CD137 (135213), APC-FasL (125239), PerCP/Cy5.5 anti-CD107a (134009), BioLegend) at 4°C for 30 min. Following staining, cells were acquired on CytoFLEX. Data analysis was performed using FlowJo software (Tree Star, BD Biosciences).
For ELISA analysis, cell culture supernatant was collected after 48 h of culture. The production of IFN-γ and TNF-α were quantified with mouse uncoated ELISA kits (88-7314-88, 88-7324-88, Thermo Scientific) according to manufacturer's instructions.

**Detection of phosphorylation of TCR signaling proteins.** For this, $CD8^+$ T cells were first stained with violet live/dead staining and seeded into the DNA disk-immobilized plates (50,000 per well) and cultured in T cell medium (no aCD28, IL-2 or IL-7) in the presence of 15 mM $MgCl_2$. The plate for co-culture was centrifuged at 500 × *g* for 2 min, followed by a co-incubation for 15 min at 37°C. Afterwards, T cells were fixed with 4% formaldehyde solution (J61899.AK, Thermo Scientific) for 15 min at room temperature to stop stimulation, and permeabilized with ice-cold methanol for 15 min on ice. The cells were then stained with mouse Alexa Fluor 647 anti-phospho-ZAP70 (PY319)/Syk (PY352) (557817, BD Biosciences), Alexa Fluor 488 anti-phospho-Erk1/2 (pT202/pY204) (612592, BD Biosciences), and PE anti-phospho-NF-κB p65 (Ser536) (5733, Cell Signaling) antibodies at 4°C for 30 min for flow cytometry analysis.

### SUPPLEMENTARY FIGURES

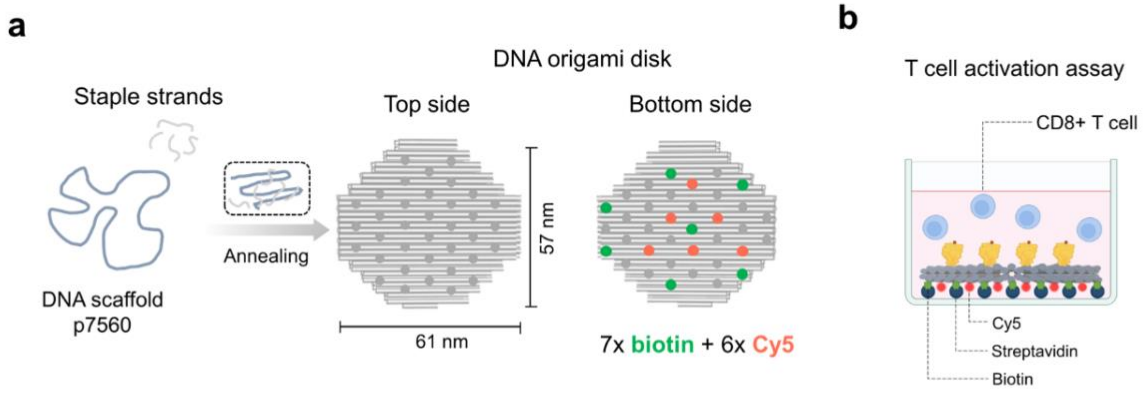


**Figure S1. Experimental setup for DNA origami-based T cell activation assay.** (a) Schematic illustration of the DNA origami disk. Each grey dot indicates an available site for ligand positioning on the top or bottom side of the DNA origami disk. The bottom side of all DNA disks was modified with integrated biotinylated strands (green dots) and Cy5 strands (red dots). (b) Schematic illustration of the T cell activation assay.

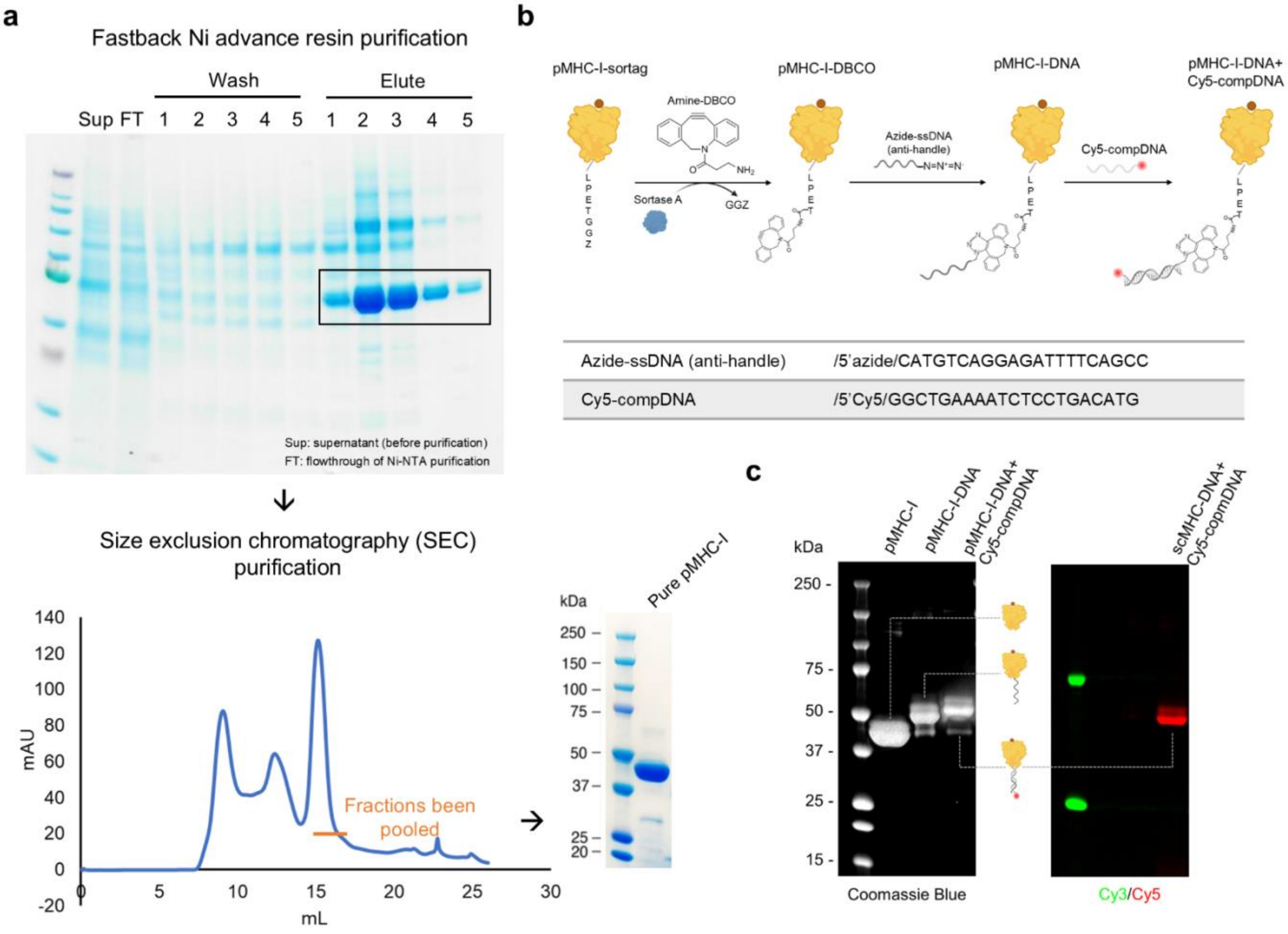


**Figure S2. Production of single-chain SIINFEK-bound MHC-I protein.** (a) His purification and size exclusion chromatography (SEC) purification of pMHC-I from Hi5 insect cell supernatant. Upper panel: SDS-PAGE analysis of eluted fractions from fastback Ni advance resin. His-tag purified fractions containing the target pMHC-I protein (indicated by black box) were further subjected to size exclusion chromatography (lower panel). Fractions corresponding to the pMHC-I monomer were pooled, yielding purified pMHC-I protein, as shown in the SDS-PAGE gel on the right. (b) Schematic illustration of the preparation of pMHC-I-DNA conjugates via sortase-mediated ligation and copper-free click chemistry. (c) SDS-PAGE analysis of pMHC-I-DNA conjugate.

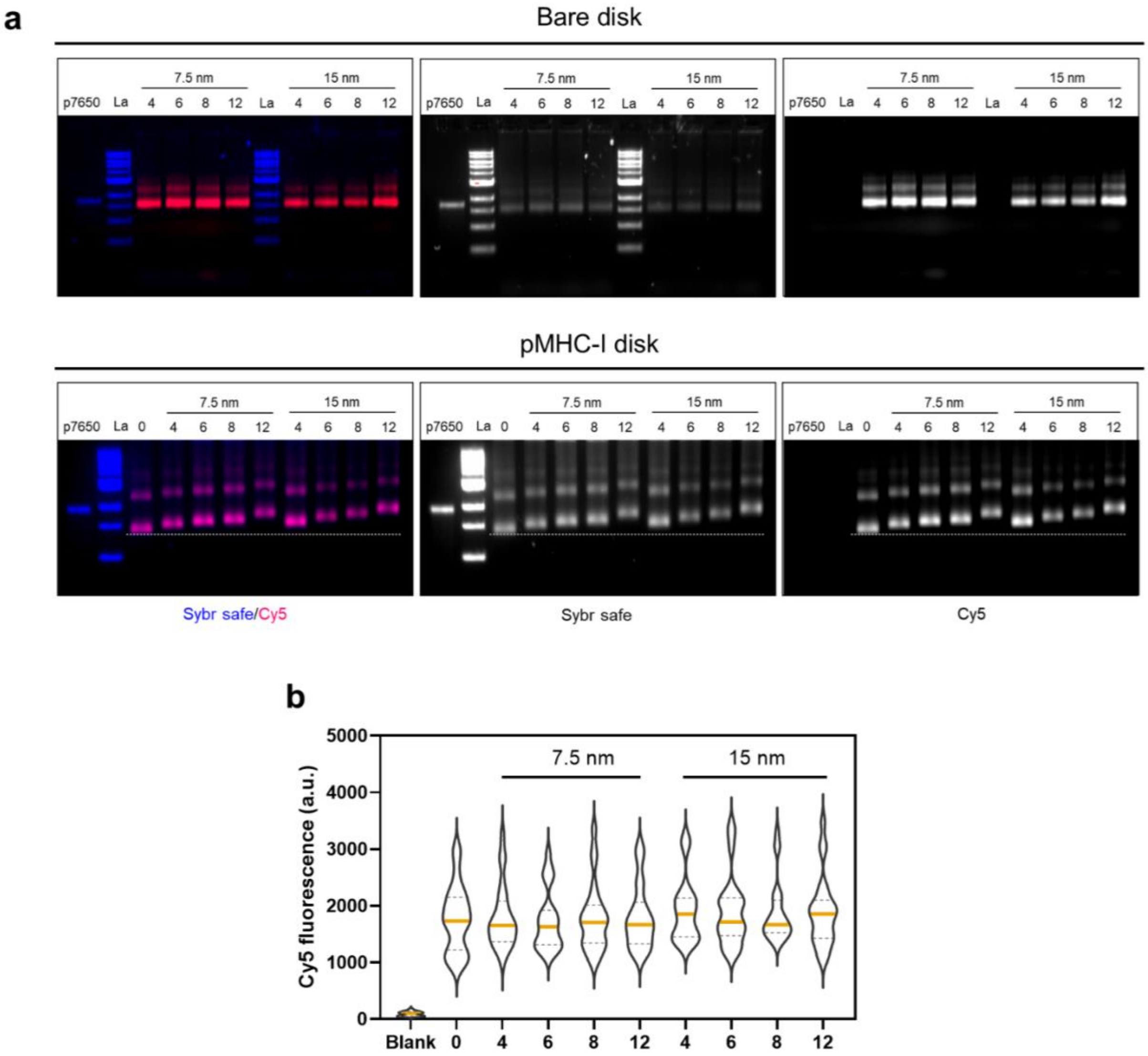


**Figure S3. Characterization of valency library of pMHC-I-decorated DNA disks.** (a) Agarose gel analysis of the DNA disk library before (1% agarose, upper panel) and after (2% agarose, lower panel) pMHC-I functionalization. SYBR Safe signal corresponds to DNA-based samples and Cy5 signal indicates folded DNA origami disk. DNA origami disks showed delayed migration after ligand attachment. (b) Representative immobilization level of DNA disks onto the cell culture plate quantified by Cy5 fluorescence intensity ($n \geqslant 30$). Blank indicates background Cy5 signal from buffer only.

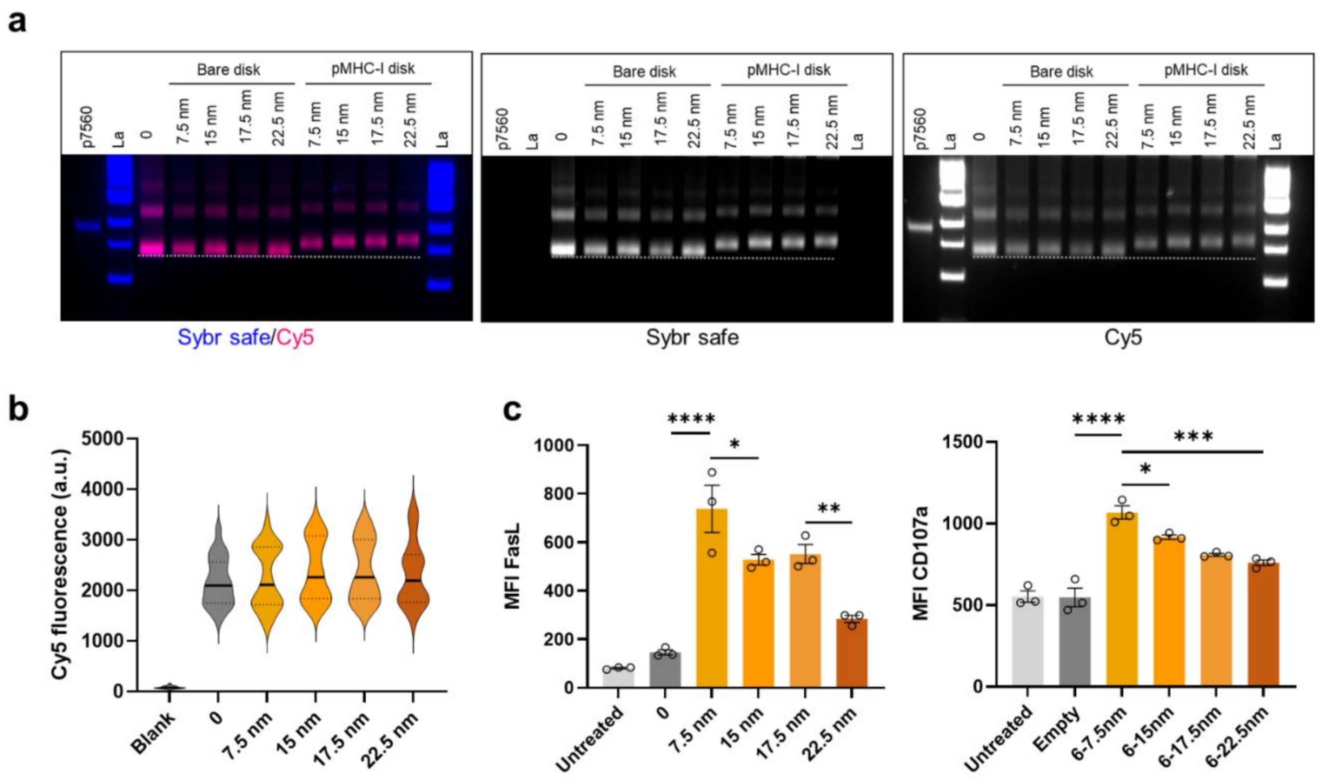


**Figure S4.** (a) Characterization of DNA disks with varying inter-ligand pMHC-I spacing by agarose gel analysis (2% agarose). SYBR Safe signal corresponds to DNA-based samples and Cy5 signal indicates folded DNA origami disk. (b) Representative immobilization level of DNA disk onto the cell culture plate quantified by Cy5 fluorescence intensity (n⩾20). Blank indicates background Cy5 signal from buffer only. (c) MFI of FasL and CD107a in $CD8^+$ T cells following 72 h stimulation by DNA origami presenting pMHC-I with varied inter-ligand spacing. Data are shown as mean ± SEM (n=3 biological replicates). Statistical significance was assessed using one-way ANOVA with Tukey's multiple comparisons test (***$p < 0.001$, ****$p < 0.0001$).

**a**

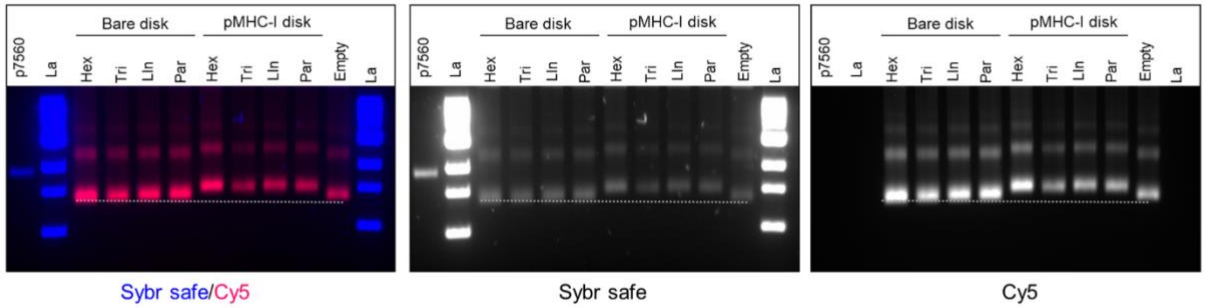


**b**

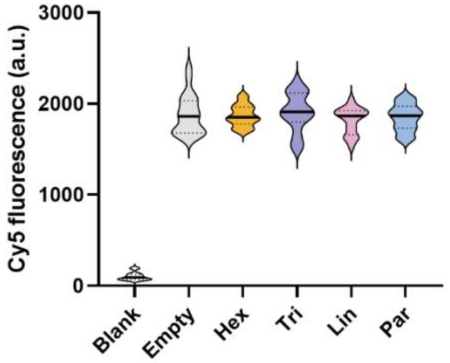


**d**

| | Hex | Tri | Lin | Par |
|---|---|---|---|---|
| Clustering area ($nm^2$) | 146.4 | 97.4 | - | 97.4 |
| pMHC-I density (per $nm^2$) | 0.041 | 0.062 | - | 0.062 |

**c**

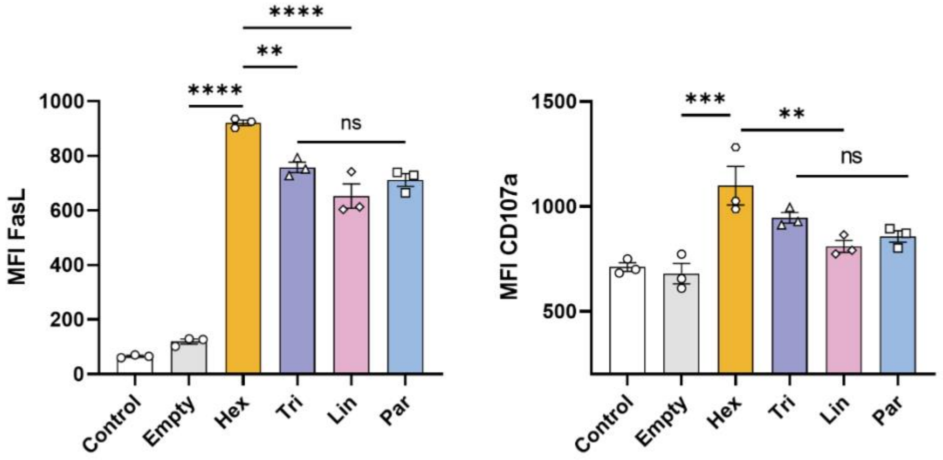


**Figure S5.** (a) Characterization of DNA disks with varying pMHC-I geometric pattern by agarose gel analysis (2% agarose). SYBR Safe signal corresponds to DNA-based samples and Cy5 signal indicates folded DNA origami disk. (b) Representative immobilization level of DNA disk onto the cell culture plate quantified by Cy5 fluorescence intensity ($n \geq 20$). Blank indicates background Cy5 signal from buffer only. (c) Expression levels of FasL and CD107a on $CD8^+$ T cells following 72 h stimulation with pMHC-I-patterned DNA disks. Data are shown as mean ± SEM (n=3 biological replicates). Statistical significance was assessed using one-way ANOVA with Tukey's multiple comparisons test (**$p < 0.01$, ***$p < 0.001$, ****$p < 0.001$). (d) Estimated average pMHC-I ligand density within individual clusters arranged in hexagonal (Hex), triangular (Tri), and parallel (Par) patterns.

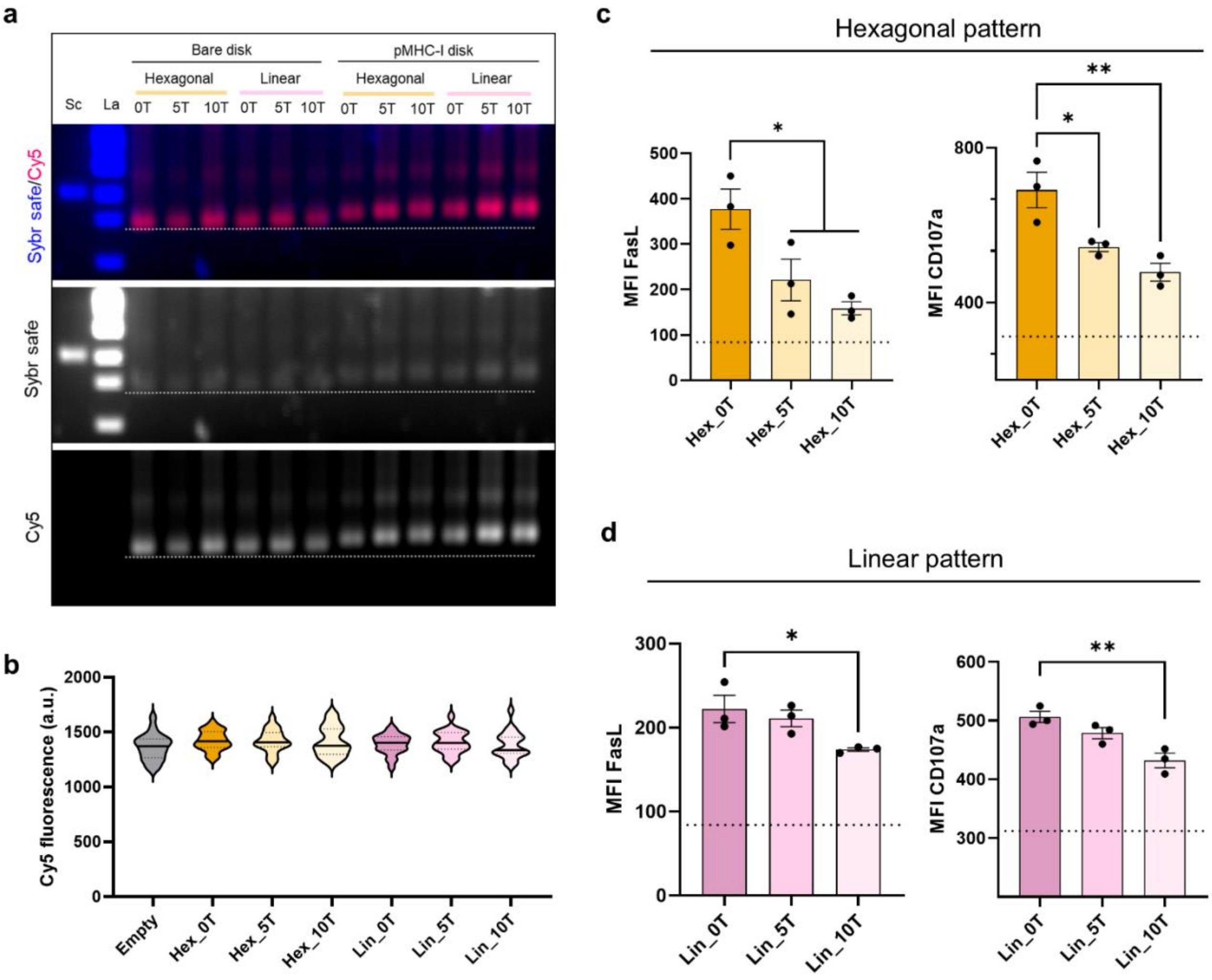


**Figure S6.** (a) Characterization of DNA disks with varying pMHC-I flexibility by agarose gel analysis (2% agarose). SYBR Safe signal corresponds to DNA-based samples and Cy5 signal indicates folded DNA origami disk. Sc: scaffold p7560. (b) Representative immobilization level of DNA disk onto the cell culture plate quantified by Cy5 fluorescence intensity (n ⩾ 20). (c) Expression levels of FasL and CD107a on CD8$^+$ T cells stimulated by hexagonally patterned pMHC-I DNA disks with varying ligand flexibilities after 72 h. (d) Expression levels of FasL and CD107a on CD8$^+$ T cells stimulated by linearly patterned pMHC-I DNA origami with varying ligand flexibilities after 72 h. The dashed line indicates the corresponding biomarker expression induced by empty DNA origami disk without pMHC-I decoration. Data are shown as mean ± SEM (n=3 biological replicates). Statistical significance was assessed using one-way ANOVA with Tukey's multiple comparisons test (*$p < 0.05$, **$p < 0.01$).

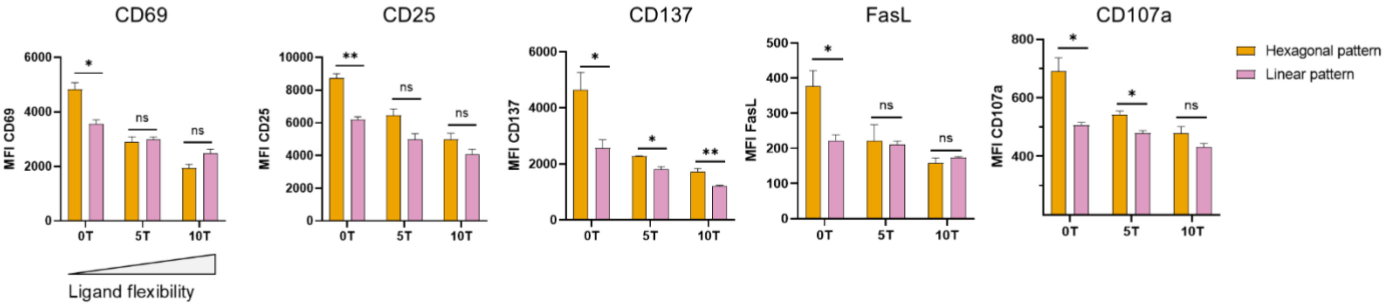


**Figure S7.** Comparative analysis of CD8+ T cell activation induced by hexagonally or linearly patterned pMHC-I DNA disks at different ligand flexibility. Data are shown as mean ± SEM (n=3 biological replicates). Statistical significance was assessed using unpaired *t* test (t (**p* < 0.05, ***p* < 0.01).

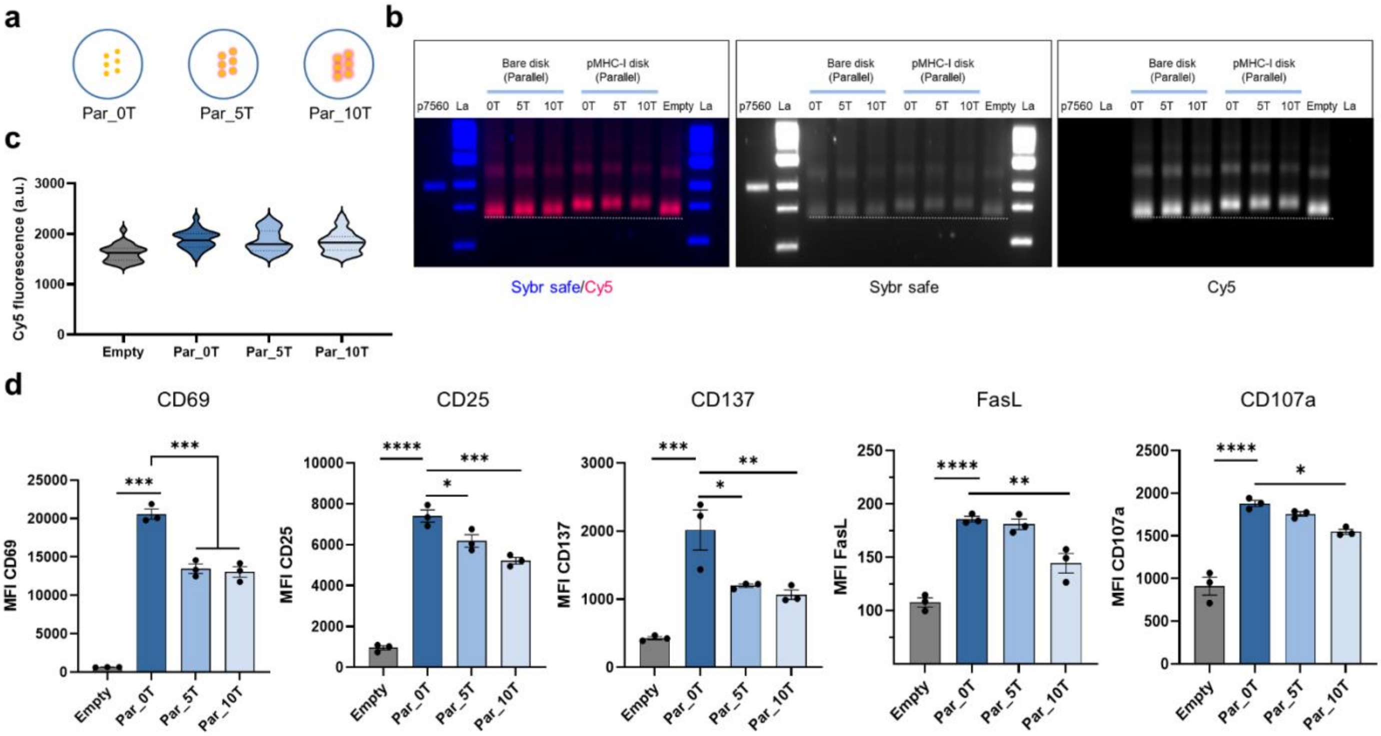


**Figure S8. Effects of pMHC-I ligand flexibility on CD8+ T cell activation, parallel pattern.** (a) Schematic illustration of DNA origami disks presenting pMHC-I with increasing ligand flexibilities arranged in a parallel pattern. (b) Characterization of DNA disks with varying pMHC-I flexibility by agarose gel analysis (2% agarose). SYBR Safe signal corresponds to DNA-based samples and Cy5 signal indicates folded DNA origami disk. (c) Representative immobilization level of DNA disk onto the cell culture plate quantified by Cy5 fluorescence intensity (n⩾20). Blank indicates background Cy5 signal from buffer only. (d) Flow cytometry analysis of CD8+ T cell activation markers following stimulation with parallelly patterned pMHC-I DNA disks of varying ligand flexibilities: CD69 (24 h), CD25 (48 h), and CD137, FasL, CD107a (72 h). Data are shown as mean ± SEM (n=3 biological replicates). Statistical significance was assessed using one-way ANOVA with Tukey's multiple comparisons test (**p* < 0.05, ***p* < 0.01, ****p* < 0.001, *****p* < 0.0001).

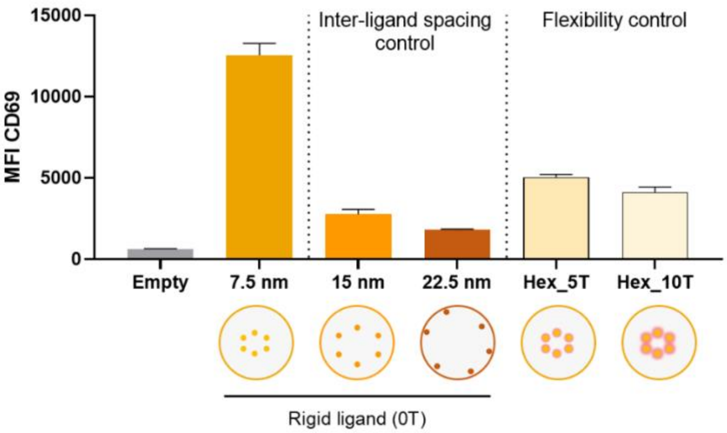


**Figure S9.** Additional experiment for comparative analysis of $CD8^+$ T cell activation potency in response to pMHC-I DNA disks with varying inter-ligand spacing and ligand flexibility. Activation was assessed by CD69 expression after 24 h stimulation. Data are shown as mean ± SD (n=3 technical replicates).

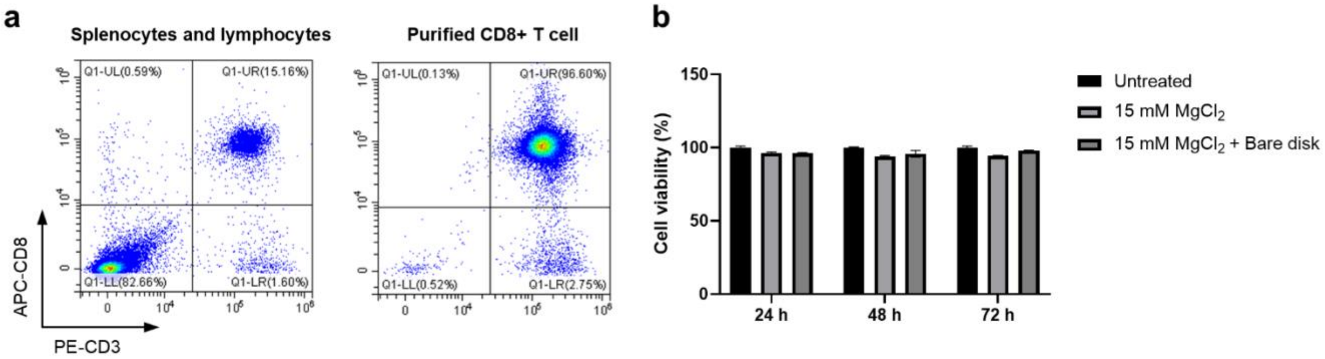


**Figure S10. Analysis of $CD8^+$ T cells purity and DNA origami disk cytotoxicity on $CD8^+$ T cells.** (a) Flow cytometry analysis of splenocytes and lymphocytes suspension (left) and purified $CD8^+$ T cells (right) before and after isolating using $CD8^+$ T cell isolation kit. (b) Cell viability of Naïve $CD8^+$ T cells treated with empty DNA origami disks and $MgCl_2$. Viability was normalized to the untreated group (set as 100%) and calculated based on $OD_{450}$ readings. Data are shown as mean ± SD (n=3 technical replicates).

### SUPPLEMENTARY TABLES

**Table S1.** Cy5-integrated staple sequences. For all Cy5-integrated DNA disk constructs, the sequences without handle extension with the corresponding numbers in Eklund *et al.*[1] were substituted with the following sequences:

| # | Name | Sequence (5'-3') |
|---|---|---|
| 5 | 5 17[77]71[87] _Cy5 | GGGACGAGTAACCGAGAACGAGCTATTT/3'Cy5/ |
| 13 | 11[140]77[150]_Cy5 | TTAAGCTCAACTCGTTCCATTACATACA/3'Cy5/ |
| 15 | 11[98]77[109]_Cy5 | CCGAACTGACGCATCTACAGACCACGGA/3'Cy5/ |
| 17 | 5[119]83[130]_Cy5 | GGCGAAACGGTCCACACCCTCTAATCAA/3'Cy5/ |
| 21 | 17[161]71[171]_Cy5 | CGCCATTGTAACAATCATTATAACAACA/3'Cy5/ |
| 26 | 17[119]71[130]_Cy5 | CAGCTTTGACCGTATTTAATCCCGACTT/3'Cy5/ |

**Table S2.** Biotin-integrated staple sequences. For all biotin-integrated DNA disk constructs, the sequences without handle extension with the corresponding numbers in Eklund *et al.*[1] were substituted with the following sequences:

| # | Name | Sequence (5'-3') |
|---|---|---|
| 14 | 3[98]85[109]_bio | GGGAAGACGTAACCGTCGAGAGAGGTTG/3'Bio/ |
| 7 | 12[44]79[46]_bio | AATACCAGTCATGGATTATTTACATCACCGAATTCATT/3'Bio/ |
| 6 | 13[119]75[130]_bio | TGCTGAAATGCGCACTGGCATAGCCGGA/3'Bio/ |
| 1 | 17[30]71[46]_bio | AAAGACAACTCGTCATTTTCTAATTTACGCTAA/3'Bio/ |
| 32 | 21[161]67[171]_bio | TAATATTTCAGGTCTTTAGACAATATTC/3'Bio/ |
| 23 | 26[107]65[109]_bio | TACGCTGAGACAACATAAATCAATCAAAAAGTAGGCAG/3'Bio/ |
| 9 | 54[175]83[172]_bio | AAAGAGTTTCGTCAC/3'Bio/ |

**Table S3.** Staple sequences for valency library of pMHC-I DNA disks. For each construct, the sequences without handle extension with the corresponding numbers in Eklund *et al.*[1] were substituted with the following sequences:

| **Design: 4 pMHC-I (inter-ligand spacing: 7.5 nm)** | | |
|---|---|---|
| **#** | **Name** | **Sequence (5'-3')** |
| 44 | 50[118]76[108] | GCATAACAGGACTAGCCTTGATCCTTAGGGCTGAAAATCTCCTGACATG |
| 48 | 45[109]72[108] | AGGGTAATGATTAGGAGCTCCAGCGGCTGAAAATCTCCTGACATG |
| 52 | 48[97]74[87] | ATAAGTTCGCAATAGGTGAGGAGTTGGCGGCTGAAAATCTCCTGACATG |
| 60 | 48[139]74[128] | TAAAGGTACTCCTTGTGGTTGTGCAAGGGGCTGAAAATCTCCTGACATG |

| **Design: 6 pMHC-I (inter-ligand spacing: 7.5 nm)** | | |
|---|---|---|
| **#** | **Name** | **Sequence (5'-3')** |
| 43 | 44[97]70[87] | AAGGCTTATTGGGCTAGATGGATGGCAAGGCTGAAAATCTCCTGACATG |
| 44 | 50[118]76[108] | GCATAACAGGACTAGCCTTGATCCTTAGGGCTGAAAATCTCCTGACATG |
| 52 | 48[97]74[87] | ATAAGTTCGCAATAGGTGAGGAGTTGGCGGCTGAAAATCTCCTGACATG |
| 60 | 48[139]74[128] | TAAAGGTACTCCTTGTGGTTGTGCAAGGGGCTGAAAATCTCCTGACATG |
| 66 | 44[139]70[129] | CATTACCATTACCTACAAACGTCTGAATGGCTGAAAATCTCCTGACATG |
| 71 | 42[118]68[107] | GCGGGAGCCGGTATAACAGAAGCCCCAAGGCTGAAAATCTCCTGACATG |

| **Design: 8 pMHC-I (inter-ligand spacing: 7.5 nm)** | | |
|---|---|---|
| **#** | **Name** | **Sequence (5'-3')** |
| 43 | 44[97]70[87] | AAGGCTTATTGGGCTAGATGGATGGCAAGGCTGAAAATCTCCTGACATG |
| 45 | 51[88]78[87] | ATTACCATAGGGAAAAACATTTCTGGCTGAAAATCTCCTGACATG |
| 50 | 54[118]80[107] | AATCACCCGGTCATGGGAAACATCGGCCGGCTGAAAATCTCCTGACATG |
| 52 | 48[97]74[87] | ATAAGTTCGCAATAGGTGAGGAGTTGGCGGCTGAAAATCTCCTGACATG |
| 53 | 52[139]78[128] | TTTGCTAACGTTGATATCCGCACAGGGCGGCTGAAAATCTCCTGACATG |
| 60 | 48[139]74[128] | TAAAGGTACTCCTTGTGGTTGTGCAAGGGGCTGAAAATCTCCTGACATG |
| 66 | 44[139]70[129] | CATTACCATTACCTACAAACGTCTGAATGGCTGAAAATCTCCTGACATG |
| 71 | 42[118]68[107] | GCGGGAGCCGGTATAACAGAAGCCCCAAGGCTGAAAATCTCCTGACATG |

| Design: 12 pMHC-I (inter-ligand spacing: 7.5 nm) | | |
| --- | --- | --- |
| # | Name | Sequence (5'-3') |
| 37 | 42[160]68[149] | TTATTACATACCACGGAACGCTAAACGTGGCTGAAAATCTCCTGACATG |
| 38 | 46[76]72[65] | TAAGCCCTAGACGGAATACATGTTTGAGGGCTGAAAATCTCCTGACATG |
| 40 | 50[160]76[149] | ATACCGATAAAATACTGCCATAAATAACGGCTGAAAATCTCCTGACATG |
| 45 | 51[88]78[87] | ATTACCATAGGGAAAAACATTTCTGGCTGAAAATCTCCTGACATG |
| 49 | 38[118]64[108] | AATGACCGGAAGCCGTCAAATAGAGTCAGGCTGAAAATCTCCTGACATG |
| 50 | 54[118]80[107] | AATCACCCGGTCATGGGAAACATCGGCCGGCTGAAAATCTCCTGACATG |
| 53 | 52[139]78[128] | TTTGCTAACGTTGATATCCGCACAGGGCGGCTGAAAATCTCCTGACATG |
| 61 | 46[160]72[149] | TTGTGTCCCAACTTCTATTACGGCAAAGGGCTGAAAATCTCCTGACATG |
| 64 | 42[76]68[65] | TGCACCCTACCGCGAGATGAAAAAATCGGGCTGAAAATCTCCTGACATG |
| 67 | 40[139]66[129] | GATAAAATTTTGCCAACAAGATCTAGCTGGCTGAAAATCTCCTGACATG |
| 69 | 40[97]66[87] | TAGAAACGTCCTGAAATCATACTTTTTGGCTGAAAATCTCCTGACATG |
| 70 | 50[76]76[65] | AAGGCCGGACAGCAGACCTGAAAACATCGGCTGAAAATCTCCTGACATG |

| Design: 4 pMHC-I (inter-ligand spacing: 15 nm) | | |
| --- | --- | --- |
| # | Name | Sequence (5'-3') |
| 43 | 44[97]70[87] | AAGGCTTATTGGGCTAGATGGATGGCAAGGCTGAAAATCTCCTGACATG |
| 45 | 51[88]78[87] | ATTACCATAGGGAAAAACATTTCTGGCTGAAAATCTCCTGACATG |
| 53 | 52[139]78[128] | TTTGCTAACGTTGATATCCGCACAGGGCGGCTGAAAATCTCCTGACATG |
| 66 | 44[139]70[129] | CATTACCATTACCTACAAACGTCTGAATGGCTGAAAATCTCCTGACATG |

| Design: 6 pMHC-I (inter-ligand spacing: 15 nm) | | |
| --- | --- | --- |
| # | Name | Sequence (5'-3') |
| 37 | 42[160]68[149] | TTATTACATACCACGGAACGCTAAACGTGGCTGAAAATCTCCTGACATG |
| 40 | 50[160]76[149] | ATACCGATAAAATACTGCCATAAATAACGGCTGAAAATCTCCTGACATG |
| 49 | 38[118]64[108] | AATGACCGGAAGCCGTCAAATAGAGTCAGGCTGAAAATCTCCTGACATG |
| 50 | 54[118]80[107] | AATCACCCGGTCATGGGAAACATCGGCCGGCTGAAAATCTCCTGACATG |
| 64 | 42[76]68[65] | TGCACCCTACCGCGAGATGAAAAAATCGGGCTGAAAATCTCCTGACATG |
| 70 | 50[76]76[65] | AAGGCCGGACAGCAGACCTGAAAACATCGGCTGAAAATCTCCTGACATG |

| Design: 8 pMHC-I (inter-ligand spacing: 15nm) | | |
|---|---|---|
| **#** | **Name** | **Sequence (5'-3')** |
| 37 | 42[160]68[149] | TTATTACATACCACGGAACGCTAAACGTGGCTGAAAATCTCCTGACATG |
| 40 | 50[160]76[149] | ATACCGATAAAATACTGCCATAAATAACGGCTGAAAATCTCCTGACATG |
| 46 | 56[97]82[87] | AGGCAGGACCAGAAAGAGCGGTCGGCCAGGCTGAAAATCTCCTGACATG |
| 47 | 33[130]62[129] | CATGCTGAATGGCTTAATTGAGTTACGCAAGACATTATGGCTGAAAATCTCCTGACATG |
| 56 | 36[97]62[87] | AGGCATTATTCTTAACCTCCGAATAAAGGGCTGAAAATCTCCTGACATG |
| 64 | 42[76]68[65] | TGCACCCTACCGCGAGATGAAAAAATCGGGCTGAAAATCTCCTGACATG |
| 68 | 56[139]82[128] | TCAGAACGGGATAGAGAGTTGAGGGTGGGGCTGAAAATCTCCTGACATG |
| 70 | 50[76]76[65] | AAGGCCGGACAGCAGACCTGAAAACATCGGCTGAAAATCTCCTGACATG |

| Design: 12 pMHC-I (inter-ligand spacing: 15 nm) | | |
|---|---|---|
| **#** | **Name** | **Sequence (5'-3')** |
| 39 | 52[186]78[171] | AAATGTCGTCTTAATTGTAAATTCGTGGAGGATGGCTGAAAATCTCCTGACATG |
| 41 | 60[99]86[87] | TACTGGTAAGTTCCAGATTTAGAAAAGGAAGGCTGAAAATCTCCTGACATG |
| 43 | 44[97]70[87] | AAGGCTTATTGGGCTAGATGGATGGCAAGGCTGAAAATCTCCTGACATG |
| 45 | 51[88]78[87] | ATTACCATAGGGAAAAACATTTCTGGCTGAAAATCTCCTGACATG |
| 47 | 33[130]62[129] | CATGCTGAATGGCTTAATTGAGTTACGCAAGACATTATGGCTGAAAATCTCCTGACATG |
| 51 | 52[55]78[44] | AGCACCGCATTTGGGTCTGAAACACGACGGCTGAAAATCTCCTGACATG |
| 53 | 52[139]78[128] | TTTGCTAACGTTGATATCCGCACAGGGCGGCTGAAAATCTCCTGACATG |
| 54 | 44[181]70[171] | GCTGACCAGGACGTTTAAATGTTCCTGTGGCTGAAAATCTCCTGACATG |
| 55 | 60[144]86[127] | AAACCCCCTGCCGTATTAAGGAACAAATAGGGTGGCTGAAAATCTCCTGACATG |
| 56 | 36[97]62[87] | AGGCATTATTCTTAACCTCCGAATAAAGGGCTGAAAATCTCCTGACATG |
| 58 | 44[55]70[44] | GATTTTTACAAAATTTTGAGTCAGAAGGGGCTGAAAATCTCCTGACATG |
| 66 | 44[139]70[129] | CATTACCATTACCTACAAACGTCTGAATGGCTGAAAATCTCCTGACATG |

**Table S5.** Staple sequences for inter-ligand spacing library of pMHC-I DNA disks. For each construct, the sequences without handle extension with the corresponding numbers in Eklund *et al.*[1] were substituted with the following sequences:

| Design: 6 pMHC-I (inter-ligand spacing: 17.5 nm) | | |
|---|---|---|
| **#** | **Name** | **Sequence (5'-3')** |
| 38 | 46[76]72[65] | TAAGCCCTAGACGGAATACATGTTTGAGGGCTGAAAATCTCCTGACATG |
| 46 | 56[97]82[87] | AGGCAGGACCAGAAAGAGCGGTCGGCCAGGCTGAAAATCTCCTGACATG |
| 47 | 33[130]62[129] | CATGCTGAATGGCTTAATTGAGTTACGCAAGACATTATGGCTGAAAATCTCCTGACATG |
| 56 | 36[97]62[87] | AGGCATTATTCTTAACCTCCGAATAAAGGGCTGAAAATCTCCTGACATG |
| 61 | 46[160]72[149] | TTGTGTCCCAACTTCTATTACGGCAAAGGGCTGAAAATCTCCTGACATG |
| 68 | 56[139]82[128] | TCAGAACGGGATAGAGAGTTGAGGGTGGGGCTGAAAATCTCCTGACATG |

| Design: 6 pMHC-I (inter-ligand spacing: 22.5 nm) | | |
|---|---|---|
| **#** | **Name** | **Sequence (5'-3')** |
| 41 | 60[99]86[87] | TACTGGTAAGTTCCAGATTTAGAAAAGGAAGGCTGAAAATCTCCTGACATG |
| 42 | 54[160]80[149] | CAGTACACCTCATAACTCACACGGAAGCGGCTGAAAATCTCCTGACATG |
| 47 | 33[130]62[129] | CATGCTGAATGGCTTAATTGAGTTACGCAAGACATTATGGCTGAAAATCTCCTGACATG |
| 51 | 52[55]78[44] | AGCACCGCATTTGGGTCTGAAACACGACGGCTGAAAATCTCCTGACATG |
| 54 | 44[181]70[171] | GCTGACCAGGACGTTTAAATGTTCCTGTGGCTGAAAATCTCCTGACATG |
| 64 | 42[76]68[65] | TGCACCCTACCGCGAGATGAAAAAATCGGGCTGAAAATCTCCTGACATG |

**Table S6.** Staple sequences for pattern library of pMHC-I DNA disks. For each construct, the sequences without handle extension with the corresponding numbers in Eklund *et al.*[1] were substituted with the following sequences:

| **Design: 6 pMHC-I (triangle pattern)** | | |
|---|---|---|
| **#** | **Name** | **Sequence (5'-3')** |
| 43 | 44[97]70[87] | AAGGCTTATTGGGCTAGATGGATGGCAAGGCTGAAAATCTCCTGACATG |
| 44 | 50[118]76[108] | GCATAACAGGACTAGCCTTGATCCTTAGGGCTGAAAATCTCCTGACATG |
| 45 | 51[88]78[87] | ATTACCATAGGGAAAAACATTTCTGGCTGAAAATCTCCTGACATG |
| 48 | 45[109]72[108] | AGGGTAATGATTAGGAGCTCCAGCGGCTGAAAATCTCCTGACATG |
| 52 | 48[97]74[87] | ATAAGTTCGCAATAGGTGAGGAGTTGGCGGCTGAAAATCTCCTGACATG |
| 60 | 48[139]74[128] | TAAAGGTACTCCTTGTGGTTGTGCAAGGGGCTGAAAATCTCCTGACATG |

| **Design: 6 pMHC-I (linear pattern)** | | |
|---|---|---|
| **#** | **Name** | **Sequence (5'-3')** |
| 44 | 50[118]76[108] | GCATAACAGGACTAGCCTTGATCCTTAGGGCTGAAAATCTCCTGACATG |
| 48 | 45[109]72[108] | AGGGTAATGATTAGGAGCTCCAGCGGCTGAAAATCTCCTGACATG |
| 49 | 38[118]64[108] | AATGACCGGAAGCCGTCAAATAGAGTCAGGCTGAAAATCTCCTGACATG |
| 50 | 54[118]80[107] | AATCACCCGGTCATGGGAAACATCGGCCGGCTGAAAATCTCCTGACATG |
| 62 | 58[118]84[107] | GCTCAGTTATAAGTGCGGTCACCCAGCAGGCTGAAAATCTCCTGACATG |
| 71 | 42[118]68[107] | GCGGGAGCCGGTATAACAGAAGCCCCAAGGCTGAAAATCTCCTGACATG |

| **Design: 6 pMHC-I (parallel pattern)** | | |
|---|---|---|
| **#** | **Name** | **Sequence (5'-3')** |
| 44 | 50[118]76[108] | GCATAACAGGACTAGCCTTGATCCTTAGGGCTGAAAATCTCCTGACATG |
| 48 | 45[109]72[108] | AGGGTAATGATTAGGAGCTCCAGCGGCTGAAAATCTCCTGACATG |
| 53 | 52[139]78[128] | TTTGCTAACGTTGATATCCGCACAGGGCGGCTGAAAATCTCCTGACATG |
| 60 | 48[139]74[128] | TAAAGGTACTCCTTGTGGTTGTGCAAGGGGCTGAAAATCTCCTGACATG |
| 66 | 44[139]70[129] | CATTACCATTACCTACAAACGTCTGAATGGCTGAAAATCTCCTGACATG |
| 71 | 42[118]68[107] | GCGGGAGCCGGTATAACAGAAGCCCCAAGGCTGAAAATCTCCTGACATG |

**Table S7.** Staple sequences for flexibility library of pMHC-I DNA disks. For each construct, the sequences without handle extension with the corresponding numbers in Eklund *et al.*[1] were substituted with the following sequences:

| Design: 6 pMHC-I (hexagonal pattern, with 5T spacer) | | |
|---|---|---|
| **#** | **Name** | **Sequence (5'-3')** |
| 43 | 44[97]70[87]_5T | AAGGCTTATTGGGCTAGATGGATGGCAATTTTTGGCTGAAAATCTCCTGACATG |
| 44 | 50[118]76[108]_5T | GCATAACAGGACTAGCCTTGATCCTTAGTTTTTGGCTGAAAATCTCCTGACATG |
| 52 | 48[97]74[87]_5T | ATAAGTTCGCAATAGGTGAGGAGTTTTTTGGCGGCTGAAAATCTCCTGACATG |
| 60 | 48[139]74[128]_5T | TAAAGGTACTCCTTGTGGTTGTGCAAGGTTTTTGGCTGAAAATCTCCTGACATG |
| 66 | 44[139]70[129]_5T | CATTACCATTACCTACAAACGTCTGAATTTTTTGGCTGAAAATCTCCTGACATG |
| 71 | 42[118]68[107]_5T | GCGGGAGCCGGTATAACAGAAGCCCCAATTTTTGGCTGAAAATCTCCTGACATG |

| Design: 6 pMHC-I (hexagonal pattern, with 10T spacer) | | |
|---|---|---|
| **#** | **Name** | **Sequence (5'-3')** |
| 43 | 44[97]70[87]_5T | AAGGCTTATTGGGCTAGATGGATGGCAATTTTTTTTTTGGCTGAAAATCTCCTGACATG |
| 44 | 50[118]76[108]_5T | GCATAACAGGACTAGCCTTGATCCTTAGTTTTTTTTTTGGCTGAAAATCTCCTGACATG |
| 52 | 48[97]74[87]_5T | ATAAGTTCGCAATAGGTGAGGAGTTTTTTTTTTTGGCGGCTGAAAATCTCCTGACATG |
| 60 | 48[139]74[128]_5T | TAAAGGTACTCCTTGTGGTTGTGCAAGGTTTTTTTTTTGGCTGAAAATCTCCTGACATG |
| 66 | 44[139]70[129]_5T | CATTACCATTACCTACAAACGTCTGAATTTTTTTTTTTGGCTGAAAATCTCCTGACATG |
| 71 | 42[118]68[107]_5T | GCGGGAGCCGGTATAACAGAAGCCCCAATTTTTTTTTTGGCTGAAAATCTCCTGACATG |

| Design: 6 pMHC-I (linear pattern, with 5T spacer) | | |
|---|---|---|
| **#** | **Name** | **Sequence (5'-3')** |
| 44 | 50[118]76[108]_5T | GCATAACAGGACTAGCCTTGATCCTTAGTTTTTGGCTGAAAATCTCCTGACATG |
| 48 | 45[109]72[108]_5T | AGGGTAATGATTAGGAGCTCCAGCTTTTTGGCTGAAAATCTCCTGACATG |
| 49 | 38[118]64[108]_5T | AATGACCGGAAGCCGTCAAATAGAGTCATTTTTGGCTGAAAATCTCCTGACATG |
| 50 | 54[118]80[107]_5T | AATCACCCGGTCATGGGAAACATCGGCCTTTTTGGCTGAAAATCTCCTGACATG |
| 62 | 58[118]84[107]_5T | GCTCAGTTATAAGTGCGGTCACCCAGCATTTTTGGCTGAAAATCTCCTGACATG |
| 71 | 42[118]68[107]_5T | GCGGGAGCCGGTATAACAGAAGCCCCAATTTTTGGCTGAAAATCTCCTGACATG |

| Design: 6 pMHC-I (linear pattern, with 10T spacer) | | |
|---|---|---|
| **#** | **Name** | **Sequence (5'-3')** |
| 44 | 50[118]76[108]_10T | GCATAACAGGACTAGCCTTGATCCTTAGTTTTTTTTTTGGCTGAAAATCTCCTGACATG |
| 48 | 45[109]72[108]_10T | AGGGTAATGATTAGGAGCTCCAGCTTTTTTTTTTGGCTGAAAATCTCCTGACATG |
| 49 | 38[118]64[108]_10T | AATGACCGGAAGCCGTCAAATAGAGTCATTTTTTTTTTGGCTGAAAATCTCCTGACATG |
| 50 | 54[118]80[107]_10T | AATCACCCGGTCATGGGAAACATCGGCCTTTTTTTTTTGGCTGAAAATCTCCTGACATG |
| 62 | 58[118]84[107]_10T | GCTCAGTTATAAGTGCGGTCACCCAGCATTTTTTTTTTGGCTGAAAATCTCCTGACATG |
| 71 | 42[118]68[107]_10T | GCGGGAGCCGGTATAACAGAAGCCCCAATTTTTTTTTTGGCTGAAAATCTCCTGACATG |

| Design: 6 pMHC-I (parallel pattern, with 5T spacer) | | |
|---|---|---|
| **#** | **Name** | **Sequence (5'-3')** |
| 44 | 50[118]76[108]_5T | GCATAACAGGACTAGCCTTGATCCTTAGTTTTTGGCTGAAAATCTCCTGACATG |
| 48 | 45[109]72[108]_5T | AGGGTAATGATTAGGAGCTCCAGCTTTTTGGCTGAAAATCTCCTGACATG |
| 53 | 52[139]78[128]_5T | TTTGCTAACGTTGATATCCGCACAGTTTTTGGCGGCTGAAAATCTCCTGACATG |
| 60 | 48[139]74[128]_5T | TAAAGGTACTCCTTGTGGTTGTGCAAGTTTTTGGGCTGAAAATCTCCTGACATG |
| 66 | 44[139]70[129]_5T | CATTACCATTACCTACAAACGTCTGAATTTTTTGGCTGAAAATCTCCTGACATG |
| 71 | 42[118]68[107]_5T | GCGGGAGCCGGTATAACAGAAGCCCCAATTTTTGGCTGAAAATCTCCTGACATG |

| Design: 6 pMHC-I (parallel pattern, with 10T spacer) | | |
|---|---|---|
| **#** | **Name** | **Sequence (5'-3')** |
| 44 | 50[118]76[108]_10T | GCATAACAGGACTAGCCTTGATCCTTAGTTTTTTTTTTGGCTGAAAATCTCCTGACATG |
| 48 | 45[109]72[108]_10T | AGGGTAATGATTAGGAGCTCCAGCTTTTTTTTTTGGCTGAAAATCTCCTGACATG |
| 53 | 52[139]78[128]_10T | TTTGCTAACGTTGATATCCGCACAGTTTTTTTTTTGGCGGCTGAAAATCTCCTGACATG |
| 60 | 48[139]74[128]_10T | TAAAGGTACTCCTTGTGGTTGTGCAAGTTTTTTTTTTGGGCTGAAAATCTCCTGACATG |
| 66 | 44[139]70[129]_10T | CATTACCATTACCTACAAACGTCTGAATTTTTTTTTTTGGCTGAAAATCTCCTGACATG |
| 71 | 42[118]68[107]_10T | GCGGGAGCCGGTATAACAGAAGCCCCAATTTTTTTTTTGGCTGAAAATCTCCTGACATG |
